# Distinct cortical-amygdala projections drive reward value encoding and retrieval

**DOI:** 10.1101/299958

**Authors:** Melissa Malvaez, Christine Shieh, Michael D. Murphy, Venuz Y. Greenfield, Kate M. Wassum

## Abstract

The value of an anticipated rewarding event is crucial information in the decision to engage in its pursuit. The networks responsible for encoding and retrieving this value are largely unknown. Using glutamate biosensors and pharmacological manipulations, we found that basolateral amygdala (BLA) glutamatergic activity tracks and mediates both the encoding and retrieval of the state-dependent incentive value of a palatable food. Projection-specific and bidirectional chemogenetic and optogenetic manipulations revealed the orbitofrontal cortex (OFC) supports the BLA in these processes. Critically, the function of ventrolateral (lOFC) and medial (mOFC) OFC→BLA projections was found to be doubly dissociable. Whereas activity in lOFC→BLA projections is necessary for and sufficient to drive encoding of a positive change in the value of a reward, mOFC→BLA projections are necessary and sufficient for retrieving this value from memory to guide its pursuit. These data reveal a new circuit for adaptive reward valuation and pursuit, indicate dissociability in the encoding and retrieval of reward memories, and provide insight into the dysfunction in these processes that characterizes myriad psychiatric diseases.

---

Prospective consideration of the outcomes of potential action choices is crucial to adaptive decision making. Chief among these considerations is the value of the anticipated rewarding events. This incentive information is state-dependent; e.g., a food outcome is more valuable when hungry than when sated. It is also learned; the value of a specific reward is encoded during its experience in a relevant motivational state ^1^. Retrieval of the previously-encoded value of an anticipated reward allows adaptive reward pursuit decisions. Dysfunction in either the value encoding or retrieval process will lead to aberrant reward pursuit and ill-informed decision making—cognitive symptoms that characterize myriad psychiatric diseases. Despite importance to understanding adaptive and maladaptive behavior, little is known of the neural circuits that support the encoding and retrieval of state-dependent reward value memories.

The basolateral amygdala (BLA) has long been known to mediate emotional learning ^2–4^ Accordingly, this structure is necessary for reward value encoding ^5–13^. But little is known of the circuitry supporting the BLA in this function. Whether the BLA participates in retrieving reward value is less clear and has been disputed ^7, 10, 11^ and its contribution, if any, to active decision making is uncertain.

## RESULTS

### BLA glutamate release tracks reward value encoding and retrieval

The BLA has intrinsic glutamatergic activity ^14–17^ and is densely innervated by glutamatergic projections from regions themselves implicated in reward learning and decision making ^18–22^. Thus, we sought to begin to fill these gaps in knowledge by using electroenzymatic biosensors to characterize BLA glutamate release during reward value encoding and retrieval. These biosensors allow sub-second, spatially-precise, sensitive, and selective measurement of neuronally-released glutamate (Fig. S1) ^23–25^. We used a behavioral paradigm that allowed us to experimentally isolate reward value encoding from the retrieval of that value and from confounding reinforcement processes (Fig. 1a) ^5, 6, 26^.

**Figure 1.**
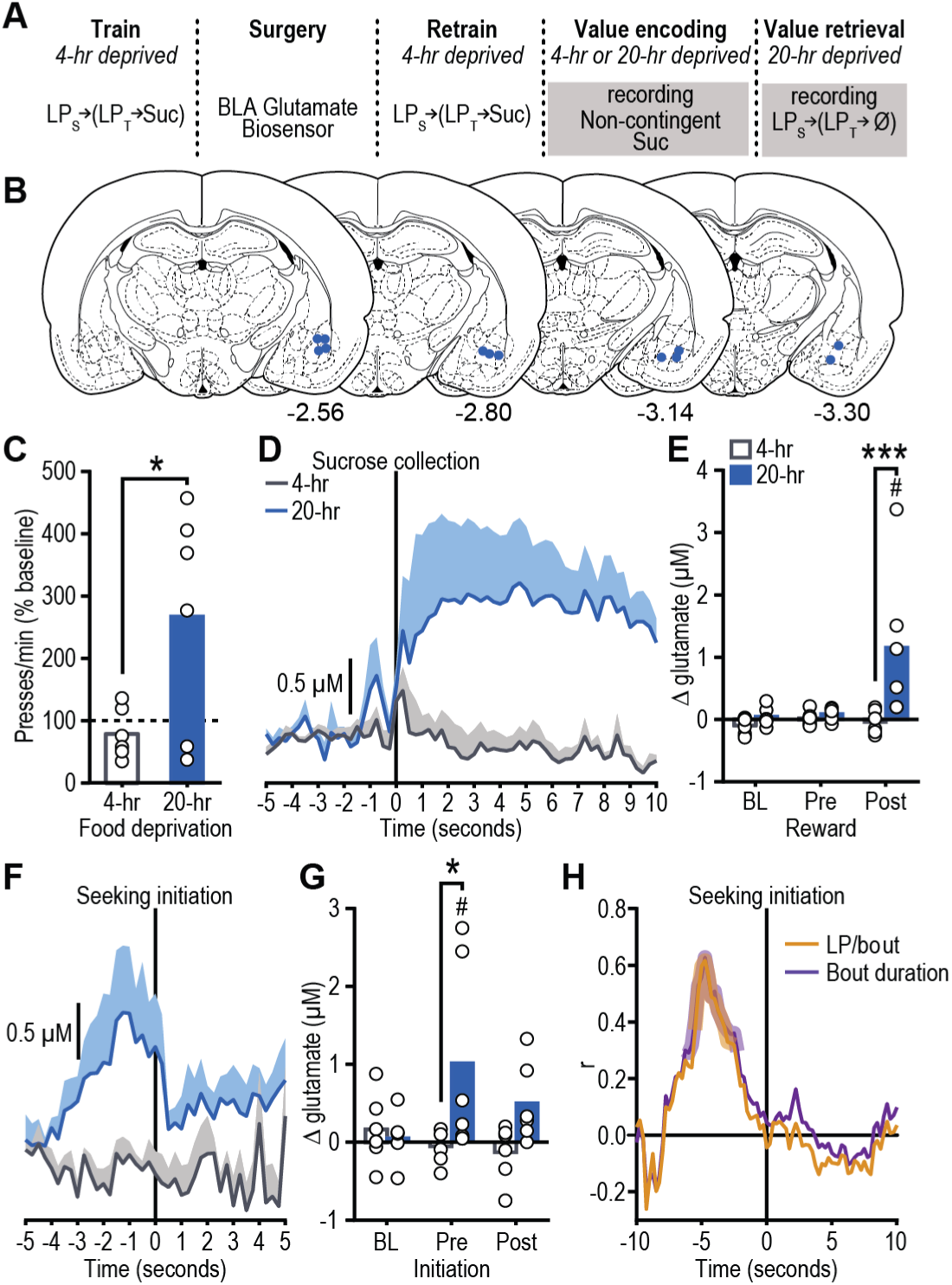
BLA glutamate release tracks reward value encoding and retrieval. (**a**) Procedure schematic (LPs, seeking lever press; LPt, taking lever press; Suc, sucrose; Ø, no sucrose delivered). (**b**) Representation of biosensor tip placements. Numbers represent anterior-posterior distance (mm) from bregma. (**c**) Reward-seeking press rate (seeking presses/min), normalized to baseline press rate (average of last-two training sessions in 4-hr food-deprived state prior to test; dashed line), during lever-pressing probe test in the hungry state for rats given prior non-contingent sucrose exposure in control sated (4-hr food-deprived; no value encoding) or hungry (20-hr deprived; value encoding opportunity) state. (**d**) Trial-averaged BLA glutamate concentration v. time trace (shading reflects between-subject s.e.m) and (**e**) quantification (mean + scatter) of average glutamate change prior to (pre) and following (post) sucrose collection/consumption (occurring at time 0 s), or equivalent baseline periods (BL) during non-contingent sucrose re-exposure in sated or hungry. (**f**) Trial-averaged BLA glutamate concentration v. time trace and (**g**) quantification of average glutamate change around bout-initiating reward-seeking presses during the lever-pressing probe test in the hungry state. (**h**) Correlation coefficient between glutamate concentration at each time point around reward-seeking bout initiation and either total seeking presses in or the duration of the subsequent bout. Shaded region indicates significant at least *P*<0.05. (*N*=6/group) \**P*<0.05, \*\**P*<0.01, between groups; ^#^*P*<0.05, relative to baseline.

Rats were trained while relatively sated (4-hr food deprived) on a self-paced, 2-lever action sequence to earn sucrose wherein pressing a ‘seeking’ lever introduced a ‘taking’ lever, a press on which retracted this lever and triggered sucrose delivery. In the sated state, the sucrose has a low value and supports a low rate of lever pressing. Once stable baseline performance was achieved, rats were re-exposed to the sucrose in either the familiar sated state or in a hungry (20-hr food-deprived) state. Because rats had never before experienced the sucrose while hungry, the latter provided an *incentive learning* opportunity to encode the high value of the sucrose in the hungry state. Re-exposure was non-contingent and conducted ‘offline’ (i.e., without the levers present) to isolate reward value encoding from reinforcement-related confounds and to prevent caching of value to the seeking and taking actions themselves. The effect of this incentive learning opportunity on rats’ reward pursuit was then tested the following day in a brief lever-pressing probe test. No rewards were delivered during this test to force the retrieval of reward value from memory and to avoid online incentive learning. Seeking presses were the primary measure because they have been shown to be selectively sensitive to learned changes in the value of an anticipated reward and relatively immune to more general motivational processes ^5, 6, 27–29^. All rats were hungry for this test, but only those rats that had previously experienced the sucrose in the hungry state escalated their reward-seeking actions (Figs. 1c, S2; *t*_10_=2.50, *p*=0.03). This result is consistent with the interpretation that the rats retrieved from memory the encoded higher value of the anticipated sucrose reward and used this information to increase their reward pursuit vigor.

BLA glutamate release was found to track reward value encoding. During re-exposure, sucrose consumption triggered a transient increase in BLA glutamate concentration, but only if a new value was being encoded (i.e., re-exposure hungry; Figs. 1d-e, S3; Time: *F*_2,20_=5.04, *P*=0.02; Deprivation: *F*_1,10_=6.67, *P*=0.03; Time × Deprivation: *F*_2,20_=4.99, *P*=0.02). This response was largest early in re-exposure (Fig. S4), when incentive learning should be the greatest. There was no BLA glutamate response detectable by the biosensor to sucrose in the absence of incentive learning either in the familiar sated state (Fig. 1d-e) or in a familiar hungry state (Fig. S5).

BLA glutamate release was also found to track reward value retrieval. In the subsequent lever-pressing test, BLA glutamate transients preceded the initiation of bouts (Table S1) of reward-seeking presses, but only if rats had prior experience with the sucrose in the hungry state and could, therefore, retrieve its current value to guide their reward pursuit actions (Figs. 1f-g, S6; Time: *F*_2,20_=1.87, *P*=0.18; Deprivation: *F*_1,10_=3.90, *P*=0.08; Time × Deprivation: *F*_2,20_=4.31, *P*=0.03). BLA glutamate transients selectively preceded the *initiation* of reward-seeking activity and did not occur prior to subsequent lever presses within a bout (Fig. S3d, S6), suggesting these signals might relate to the considerations driving reward pursuit. This was further supported by evidence that the magnitude of pre-bout-initiation BLA glutamate release positively correlated on a trial × trial basis with the number of seeking presses in and duration of the subsequent bout (presses: r_88_=0.23, *p*=0.03, duration: r_88_=0.21, *p*=0.05); longer bouts of reward seeking were preceded by larger amplitude glutamate transients. In the group that received incentive learning, glutamate release magnitude significantly predicted future reward-seeking activity in the seconds prior to, but not following the initiation of reward seeking (Fig. 1h).

### BLA glutamate receptor activity is necessary for reward value encoding and retrieval

We next assessed whether BLA glutamate activity is necessary for the encoding and/or retrieval of reward value by blocking BLA glutamate receptors during either sucrose re-exposure (encoding) or during the post-re-exposure lever-pressing test (retrieval) (Fig. 2). Following training in the sated state, all rats were provided the incentive learning opportunity (sucrose re-exposure hungry; Fig. 2a). Inactivation of neither NMDA, with infenprodil ^7, 30^, nor AMPA, with NBQX ^31^, receptors in the BLA altered food-port checking behavior (Fig. 2c; *F*_2,23_=0.81, *P*=0.46) or sucrose palatability responses (Fig. 2d; *F*_2,21_=0.12, *P*=0.88) during re-exposure. Inactivation of BLA NMDA, but not AMPA receptors did, however, prevent the subsequent upshift in reward seeking that would have otherwise occurred when animals were tested in the hungry state drug-free the next day (Figs. 2e, S7; *F*_2,23_=4.48, *P*=0.03), indicating that BLA NMDA receptors are necessary for assigning positive value to a reward. All rats were then given the incentive learning opportunity drug-free, and were tested again for their lever pressing in the hungry state on drug (Fig. 2f). In this case, either BLA AMPA or NMDA receptor inactivation prevented the increase in value-guided reward seeking that should have occurred following incentive learning (Fig. 2g, S8; *F*_2,19_=7.22, *P*=0.005). Therefore, BLA glutamate signaling tracks and is necessary for both reward value encoding and value-guided reward pursuit.

**Figure 2.**
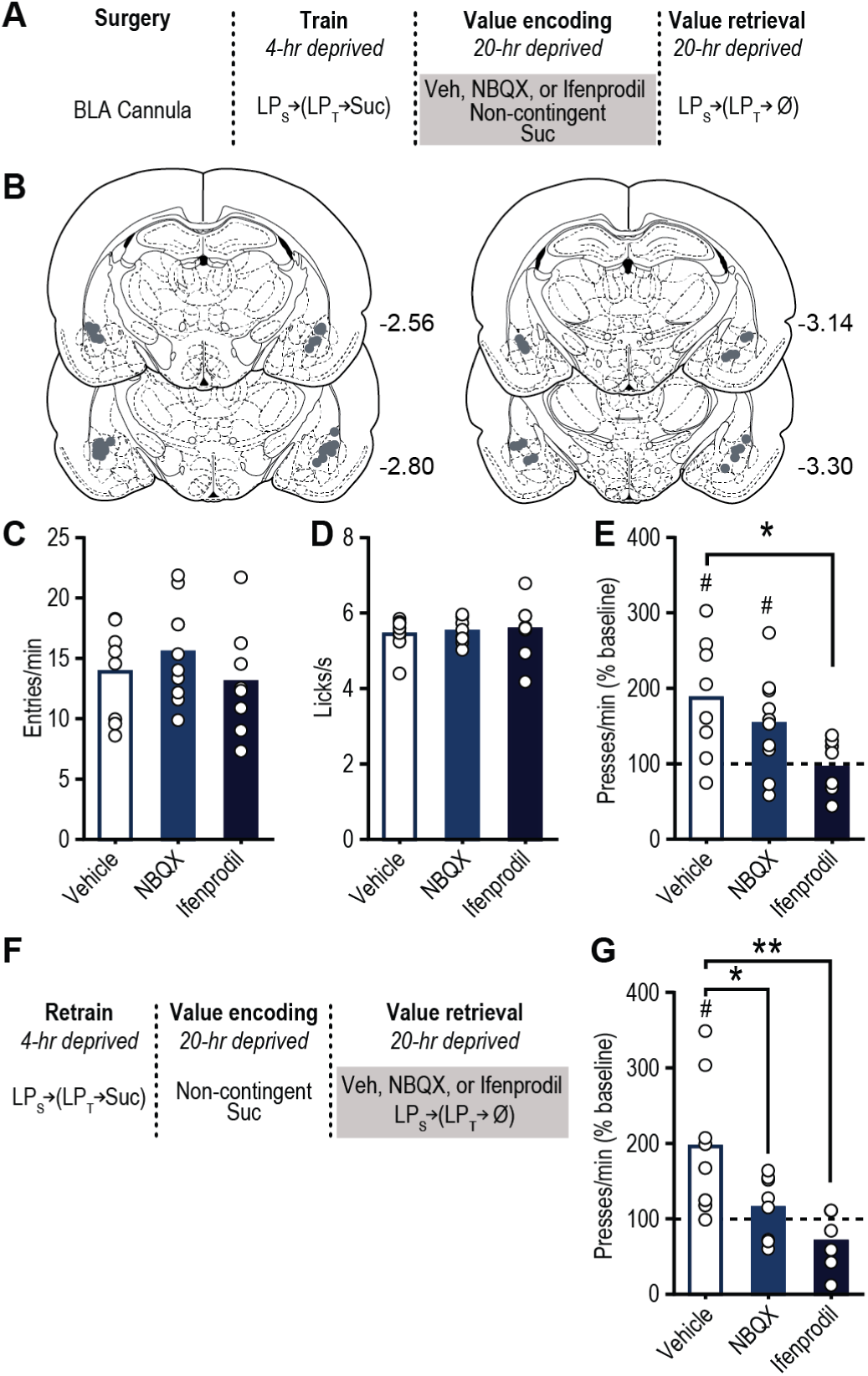
BLA glutamate receptor activity is necessary for reward value encoding and retrieval. (**a**) Procedure schematic (LPs, seeking lever press; LPt, taking lever press; Suc, sucrose; Ø, no sucrose delivery; Veh, Vehicle; NBQX, AMPA antagonist; Ifenprodil, NMDA antagonist). (**b**) Microinfusion injector tip placements. Numbers represent anterior-posterior distance (mm) from bregma. (**c**) Food-port entry rate (entries/min) and (**d**) palatability responses (lick frequency; licks/s) during non-contingent sucrose re-exposure in hungry state (20-hr deprived; value encoding opportunity) following intra-BLA infusion of Vehicle (*N*=8), AMPA (*N*=10), or NMDA (*N*=9) antagonist. (**e**) Reward-seeking press rate (seeking presses/min), relative to baseline press rate (dashed line), during drug-free, lever-pressing probe test in hungry state. (**f**) Procedure schematic. (**g**) Following off-drug sucrose re-exposure in hungry state, reward-seeking press rate, relative to baseline, during the on-drug (intra-BLA Vehicle (*N*=8), AMPA (*N*=8), or NMDA (*N*=7) antagonist) lever-pressing probe test in the hungry state. \**P*<0.05, \*\**P*<0.01 between groups; ^#^*P*<0.05 relative to baseline. Data presented as mean + scatter.

### Distinct OFC→BLA projections are necessary for reward value encoding and retrieval

An excitatory input to the BLA might facilitate its function in reward value encoding and retrieval. The orbitofrontal cortex (OFC) is a prime candidate for this because it sends dense glutamatergic innervation to the BLA ^18–22^ and is itself implicated in reward processing and decision making ^32–41^, including incentive learning ^42^. So we next used a chemogenetic approach and the same behavioral task to ask whether OFC→BLA projections are necessary for reward value encoding and/or retrieval (Fig. 3). The lateral (lOFC) and medial (mOFC) OFC subdivisions are anatomically and functionally distinct ^43–47^. We identified projections to the BLA from both the mOFC and lOFC (Fig. S9a-b). Therefore, we assessed the function of both lOFC→BLA and mOFC→BLA projections in reward value encoding and retrieval.

**Figure 3.**
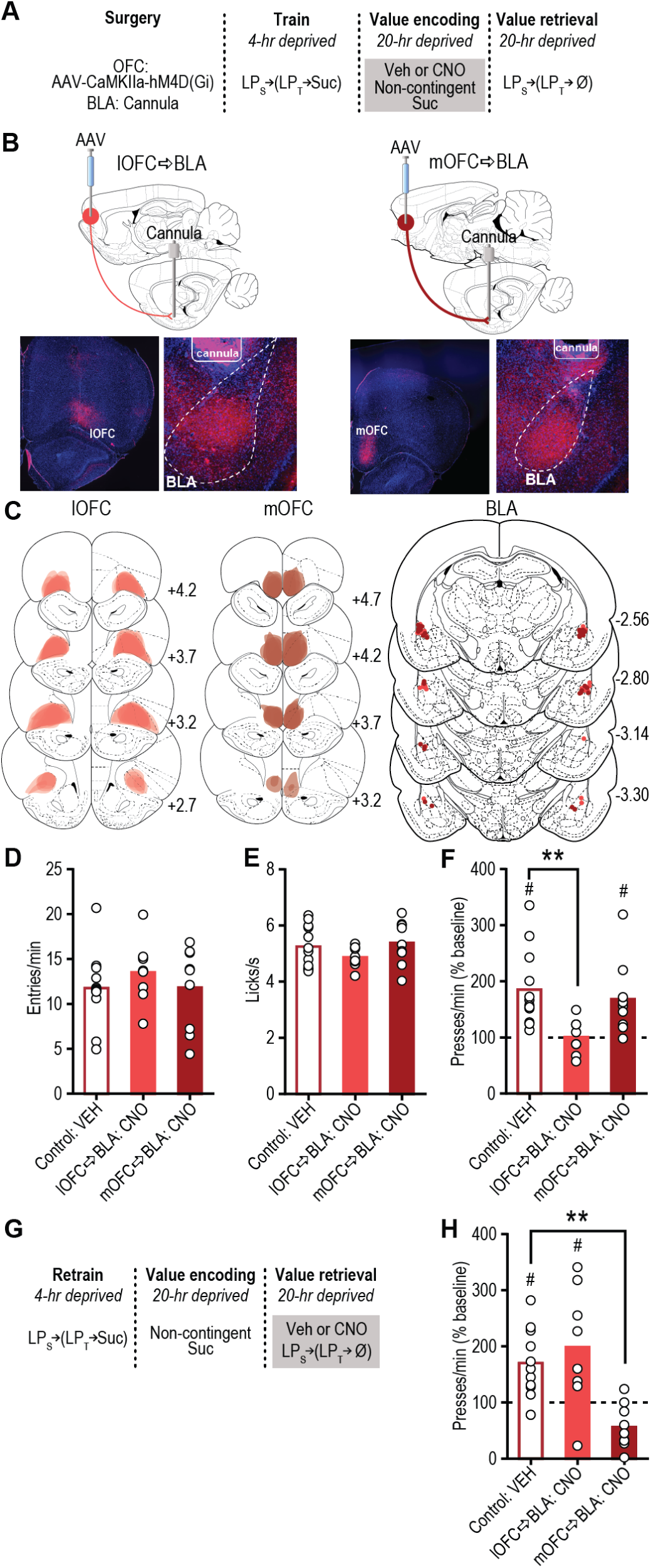
lOFC→BLA and mOFC→BLA projections are necessary for reward value encoding and retrieval, respectively. (**a**) Procedure schematic (LPs, seeking lever press; LPt, taking lever press; Suc, sucrose; Ø, no sucrose delivery; Veh, Vehicle; CNO, Clozapine N-oxide). (**b**) *Top*, schematic of chemogenetic approach for inactivation of lOFC (*left*) or mOFC (*right*) terminals in the BLA. *Bottom*, representative immunofluorescent images of HA-tagged hM4D(Gi) expression in lOFC (*left*) or mOFC (*right*) and cannula above terminal expression in the BLA. (**c**) Schematic representation of hM4D(Gi) expression in lOFC or mOFC and placement of microinfusion injector tips in the BLA for all subjects. Numbers represent anterior-posterior distance (mm) from bregma. (**d**) Food-port entry rate (entries/min) and (**e**) palatability responses (lick frequency, licks/s) during non-contingent sucrose re-exposure in hungry state (20-hr deprived; value encoding opportunity) following intra-BLA infusion of Vehicle (*N*=12, ½ mOFC hM4D(Gi), ½ lOFC hM4D(Gi)) or CNO (1mM/0.5µl; lOFC→BLA:CNO, *N*=8; mOFC→BLA:CNO, *N*=9). (**f**) Reward-seeking press rate (seeking presses/min), relative to baseline press rate (dashed line), during drug-free, lever-pressing probe test in hungry state. (**g**) Procedure schematic. (**h**) Following off-drug sucrose re-exposure in hungry state, reward-seeking press rate, relative to baseline, during the on-drug test (intra-BLA Vehicle (*N*=11) or CNO (lOFC→BLA:CNO, *N*=8; mOFC→BLA:CNO, *N*=9)). \*\**P*<0.01, between groups; ^#^*P*<0.05 relative to baseline. Data presented as mean + scatter.

Rats expressing the inhibitory designer receptor *human M4 muscarinic receptor* (hM4D(Gi)) in excitatory cells of either the lOFC or mOFC showed robust expression in terminals in the BLA in the vicinity of implanted guide cannula (Fig. 3b-c; see also Fig. S9c-d). Clozapine N-oxide (CNO; 1mM/0.5μl) was infused into the BLA to inactivate these terminals (Fig. S10) ^48^ during the sucrose re-exposure, incentive learning opportunity and lever pressing was assessed the following day drug-free (Fig. 3a). Neither manipulation altered food-port checking behavior (Fig. 3d; *F*_2,26_=0.54, *P*=0.59) or sucrose palatability responses (Fig. 3e; *F*_2,26_=1.33, *P*=0.28) online during the re-exposure. Inhibition of lOFC, but not mOFC terminals in the BLA did, however, prevent the subsequent upshift in reward seeking that would have otherwise occurred (Figs. 3f, S11; *F*_2,26_=5.06, *P*=0.014). These data suggest that activity in lOFC→BLA, but not mOFC→BLA projections is necessary for encoding the positive value of a rewarding event.

To determine whether OFC→BLA projections are necessary for reward value retrieval, we allowed all the rats to encode the sucrose’s high value in the hungry state drug-free and then evaluated their lever pressing in the hungry state following intra-BLA vehicle or CNO infusion (Fig. 3g). In this case, inhibition of mOFC, but not lOFC terminals in the BLA attenuated reward-seeking activity (Figs. 3h, S12; *F*_2,25_=9.81, *P*=0.0007), without altering performance of other indices of motivated behavior (Fig. S12). Inactivation of mOFC→BLA projections was without effect if reward value was not being retrieved from memory, either because it had not been learned or because it was observable to the subject and could, therefore, be held in working memory at test (Fig. S13). These data indicate the necessity of activity in mOFC→BLA, but not lOFC→BLA projections in retrieving the value of an anticipated reward. Thus, lOFC→BLA projections are necessary for encoding reward value, but their activity is not necessary to retrieve this information. Whereas mOFC→BLA projections are not necessary for encoding reward value, but are required to retrieve this information. Secondarily, this double dissociation indicates the behavioral effects are not simply due to off-target effects of CNO itself in the absence of hM4D(Gi), which would cause uniform behavioral effects regardless of hM4D(Gi) subregion.

### Optical stimulation of lOFC→BLA, but not mOFC→BLA projections is sufficient to instantiate value to a specific reward

That lOFC→BLA projections were necessary for positive reward value encoding, suggests that activity in these projections might drive such encoding. To test this, we optically stimulated lOFC terminals in the BLA (Fig. S10) concurrent with sucrose experience under conditions in which incentive learning would not normally occur: a familiar sated state (Fig. 4a). In a separate group, we stimulated mOFC terminals in the BLA. We restricted optical stimulation (473 nm, 20Hz, 10mW, 5 s) to the time of sucrose consumption during non-contingent exposure to match the timing of BLA glutamate release detected during incentive learning (Fig. 1d). Rats expressing the excitatory opsin *channelrhodopsin-2* (ChR2) in excitatory cells of either the lOFC or mOFC showed robust expression in terminals in the BLA in the vicinity of implanted optical fibers (Fig. 4b-c; see also Fig. S9e-f). Stimulation of lOFC terminals in the BLA concurrent with reward consumption in the familiar sated state did not alter food-port checking behavior (Fig. 4d; *t*_16_=0.20, *p*=0.84) or sucrose palatability responses (Fig. 4e; *t*_16_=0.25, *p*=0.80) online. But it did cause a dramatic increase in reward-seeking presses in the test conducted in that same sated state manipulation-free the following day (Figs. 4f, S14; *F*_2,24_=9.25, *P*=0.001), mimicking the effect of hunger-induced incentive learning (Fig. S15). This did not occur under otherwise identical circumstances with stimulation paired with a task-irrelevant rewarding event (food pellet), ruling out the confounding possibility of enhanced context salience or other factors unrelated to motivation to obtain the specific anticipated sucrose reward (Fig. 4f). lOFC→BLA stimulation also amplified normal, hunger-induced incentive learning (Fig. S15). Identical stimulation of mOFC terminals in the BLA had no effect on online food-port checking behavior (Fig. 4d; *t*_10_=0.49, *p*=0.64) or sucrose palatability responses (Fig. 4e; *t*_10_=0.07, *p*=0.95), or on subsequent reward-seeking presses (Figs. 4f, S14; *t*_10_=1.17, *p*=0.27). Thus, activity in lOFC→BLA, but not mOFC→BLA projections is sufficient to instantiate value to a rewarding event, and, thereby, drive escalation of its pursuit.

**Figure 4.**
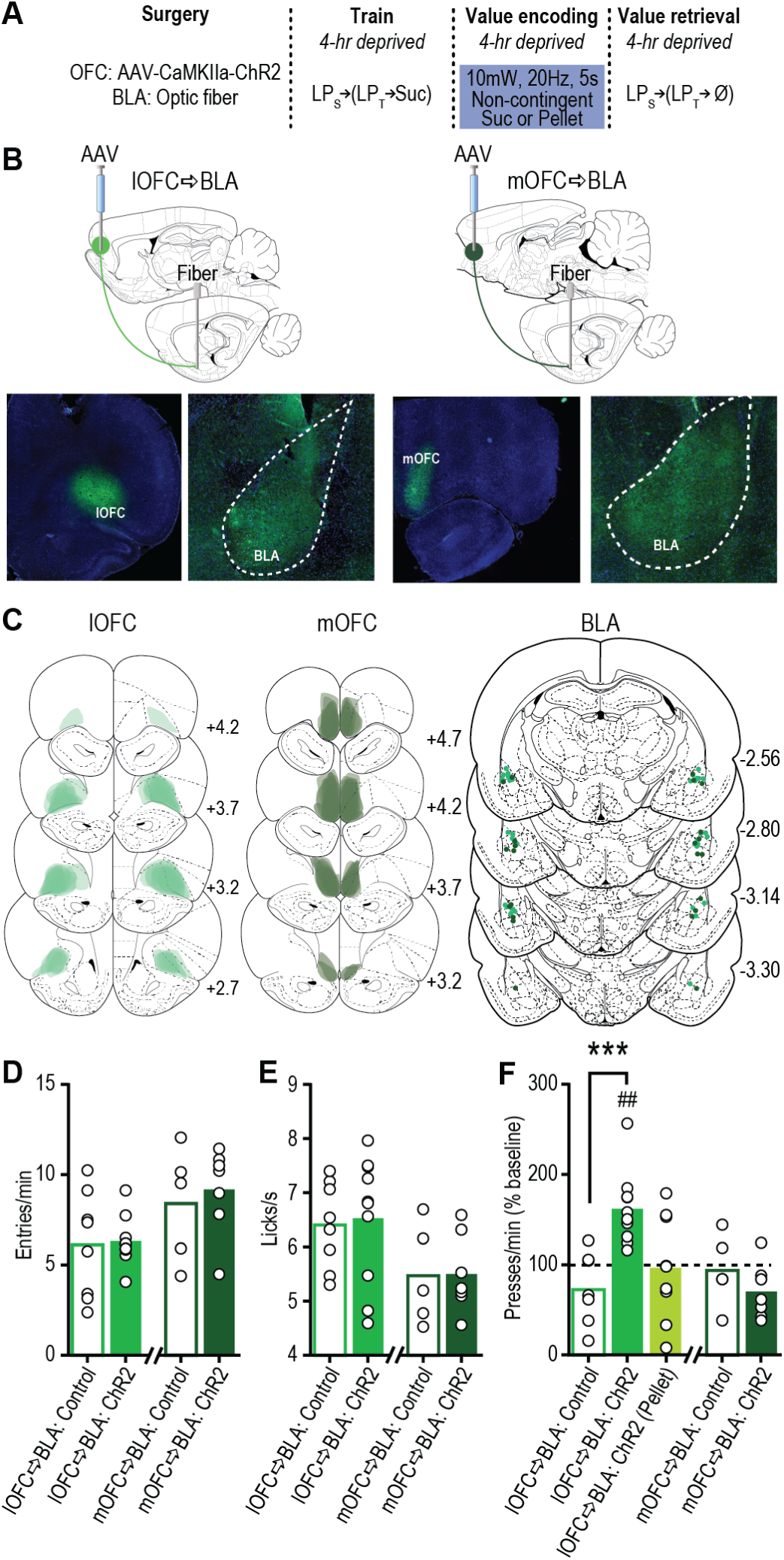
Optical stimulation of lOFC terminals in BLA concurrent with reward experience is sufficient to drive positive value assignment. (**a**) Procedure schematic (LPs, seeking lever press; LPt, taking lever press; Suc, sucrose; Ø, no sucrose delivery). (**b**) *Top*, schematic of optogenetic approach for stimulation of lOFC (*left*) or mOFC (*right*) terminals in the BLA. *Bottom*, representative fluorescent images of ChR2-eYFP expression in lOFC (*left*) or mOFC (*right*) and BLA terminal field. (**c**) Schematic representation of ChR2 expression in lOFC or mOFC and placement of optical fiber tips in BLA for all subjects. Numbers represent anterior-posterior distance (mm) from bregma. (**d**) Food-port entry rate (entries/min) and (**e**) palatability responses (lick frequency, licks/s) during non-contingent sucrose re-exposure in control 4-hr food-deprived sated state. Light (10mW, 20Hz, 5 s) was delivered concurrent with each sucrose collection. Control groups consisted of half eYFP-only + 473 nm and half ChR2 + 589 nm light delivery. (**f**) Reward-seeking press rate (seeking presses/min), relative to baseline press rate (dashed line), during manipulation-free, lever-pressing probe test in sated state. (Pellet) refers to the control condition of optical stimulation of lOFC terminals in BLA paired with stimulation of a task-irrelevant food pellet rather than sucrose. lOFC→BLA:Control, *N*=8; lOFC→BLA:ChR2, *N*=10; lOFC→BLA:ChR2 (Pellet), *N*=9; mOFC→BLA:Control, *N*=5; mOFC→BLA:ChR2, *N*=7. \*\*\**P*<0.001, between groups; ^##^*P*<0.01, relative to baseline. Data presented as mean + scatter.

### Optical stimulation of mOFC→BLA, but not lOFC→BLA projections is sufficient to facilitate reward value retrieval

That mOFC→BLA projections were necessary for reward value retrieval suggests activity in these projections might facilitate the retrieval of the value of an anticipated reward. If this is true, then optically stimulating mOFC→BLA projections during lever pressing should enhance reward seeking following an incentive learning opportunity that would not itself support an upshift in reward pursuit. To test this, we expressed ChR2 in the mOFC and, following sucrose re-exposure in a moderate, 8hr food-deprived hunger state, optically stimulated mOFC terminals in the BLA during a lever-pressing test conducted in that same 8-hr moderate hunger state (Fig. 5a-c). A separate group received stimulation of lOFC terminals in the BLA. In controls, sucrose exposure in the 8-hr food-deprived state was not sufficient to drive an increase in reward pursuit when tested the following day in this state, confirming subthreshold incentive learning (Fig. 5d). Stimulation of mOFC terminals in the BLA (473 nm, 20Hz, 10mW, 3 s, once/min) promoted reward-seeking activity under these conditions (Figs. 5d, S16; *t*_15_=3.62, *p*=0.003). Stimulation did not increase reward seeking when tested in the well-learned low-value satiety state, or following effective incentive learning in the high-value hungry state (Fig. S17). mOFC→BLA stimulation was also without effect under otherwise identical circumstances in the absence of the subthreshold incentive learning opportunity (Fig. 5e-f, S18; *t*_8_=0.67, *p*=0.52), isolating its effect to reward value retrieval. Stimulation of lOFC→BLA projections during the reward-seeking test had no effect (Figs. 5d, S16; *t*_11_=0.737, *p*=0.72). These data indicate that activity in mOFC→BLA, but not lOFC→BLA projections is sufficient to facilitate state-dependent reward value retrieval.

**Figure 5.**
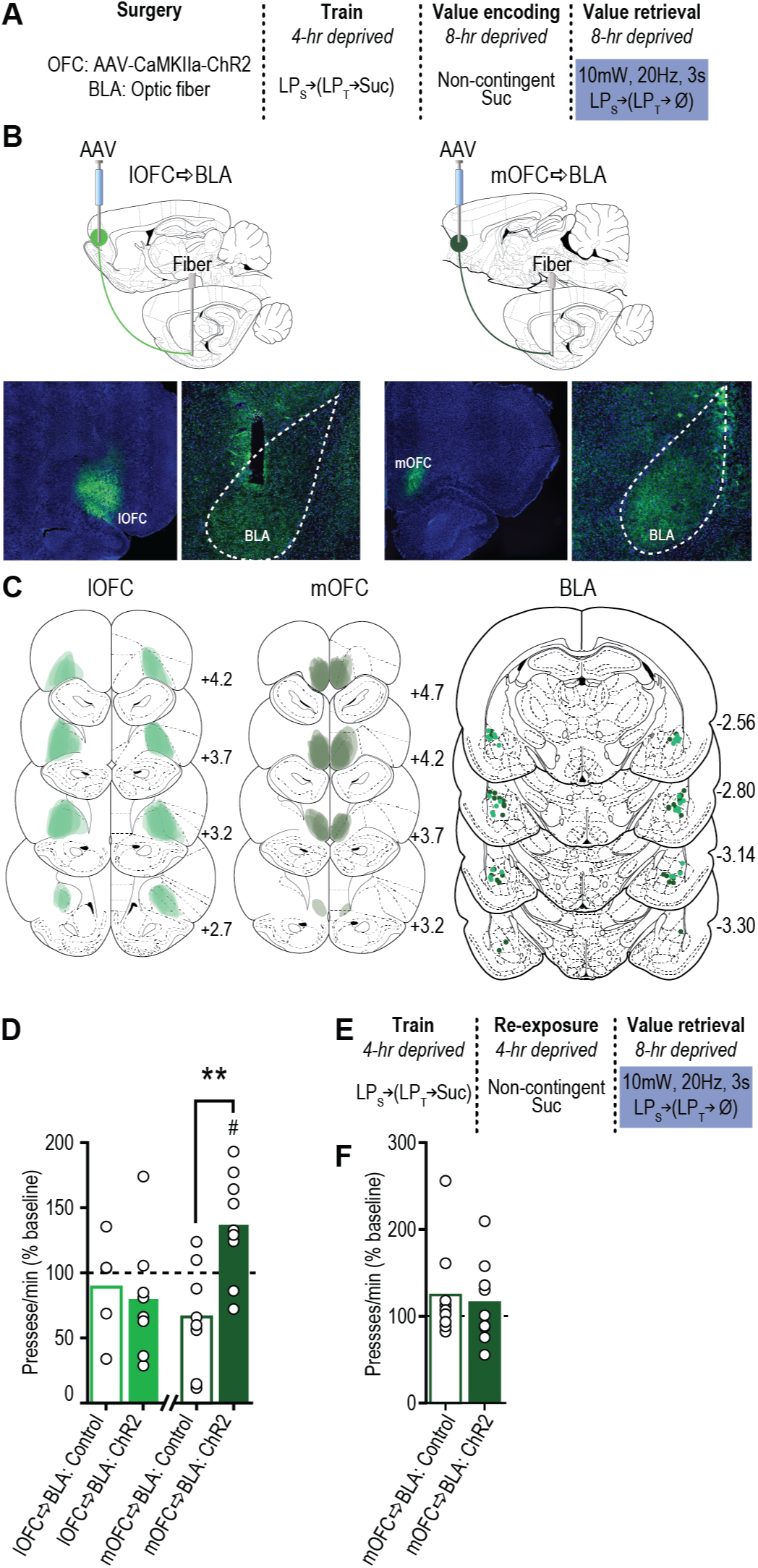
Optical stimulation of mOFC→BLA projections is sufficient to enhance reward value retrieval. (**a**) Procedure schematic (LPs, seeking lever press; LPt, taking lever press; Suc, sucrose; Ø, no sucrose delivered). (**b**) *Left*, schematic of optogenetic approach for stimulation of lOFC (*top*) or mOFC (*bottom*) terminals in the BLA. *Right*, representative fluorescent images of ChR2-eYFP expression in lOFC (*top*) or mOFC (*bottom*) and BLA terminal field. (**c**) Schematic representation of ChR2 expression in lOFC or mOFC and placement of fiber tips in BLA for all subjects. Numbers represent anterior-posterior distance (mm) from bregma. (**d**) Reward-seeking press rate (seeking presses/min), relative to baseline press rate (dashed line), during lever-pressing probe test in moderate (8-hr food-deprived) hunger state following sucrose re-exposure in 8-hr food-deprived state (a sub-threshold incentive learning opportunity). Light (10mW, 20Hz, 3 s, once/min) was delivered during this test. Control groups consisted of half eYFP-only + 473 nm and half ChR2 + 589 nm light delivery. (**e**) Procedure schematic for a separate group of all ChR2-expressing subjects for which the sub-threshold incentive learning was omitted. (**f**) Reward-seeking press rate, relative to baseline press rate, during lever-pressing probe test in moderate (8-hr food-deprived) hunger state following sucrose re-exposure in 4-hr food-deprived state. Light (10mW, 20Hz, 3 s, once/min) was delivered during this test. Within-subject control consisted of identical delivery of 589 nm light during lever pressing test (test order counterbalanced). lOFC→BLA:Control, *N*=5; lOFC→BLA:ChR2, *N*=8; mOFC→BLA:Control, *N*=8; mOFC→BLA:ChR2, *N*=9; mOFC→BLA(no incentive learning), *N*=9 \*\**P*<0.01, between groups; ^#^*P*<0.05, relative to baseline. Data presented as mean + scatter.

## DISCUSSION

These data provide evidence for the BLA as a crucial locus for not only learning about the value of a rewarding event, but also for retrieving this information to guide adaptive reward pursuit, revealing it as a critical contributor to value-based decision making. These value encoding and retrieval functions were found to be supported via doubly dissociable contributions of excitatory input from the lOFC and mOFC. Whereas lOFC→BLA projection activity is necessary for and sufficient to drive encoding of a reward’s positive value, it does not mediate the retrieval of that state-dependent reward value memory. Conversely, activity in mOFC→BLA projections does not mediate reward value encoding, but it is necessary and sufficient for retrieving an anticipated reward’s value from memory to guide reward pursuit decisions.

BLA glutamate activity was found to track and mediate both reward value encoding and retrieval. The necessity of BLA NMDA receptors for incentive learning is consistent with the long-standing crucial role for these receptors in BLA synaptic plasticity ^49–51^ and in establishing long-term, BLA-dependent memories ^52, 53^. Following a learning event, AMPA receptors are trafficked to the membrane ^54^ and such trafficking can regulate the expression of NMDA-dependent synaptic plasticity in the BLA ^55^. Consistent with this, we found that value-guided reward seeking requires BLA AMPA receptor activation, as well as NMDA receptors.

This role for BLA in reward value encoding and retrieval accords well with previous evidence of BLA necessity for reward value learning ^5, 7, 8, 10, 12^, but differs from data demonstrating the BLA is not required for value retrieval following sensory-specific satiety devaluation ^7, 10, 11^. In these latter experiments, the value shift was negative, temporary, and occurred immediately prior to test. Our value learning was positive, permanent, and occurred at least 24 hr prior to test. We suggest, therefore, that the BLA facilitates the encoding and retrieval of long-term, need-state-dependent reward value memories, and, as such, is a critical contributor to value-based decision making. This interpretation is consistent with evidence from humans and non-human primates that BLA activity can encode value ^56^, prospectively reflect goal plans ^57^, and predict behavioral choices ^58^, and with evidence of temporally-specific BLA inactivation disrupting choice behavior ^59^. Via direct projections, the OFC was found to support the BLA in its function in reward value. This is consistent with evidence of broad OFC encoding of reward value across species ^60–69^ and with evidence of cooperative OFC and amygdala function in reward learning and choice in rodents and primates ^70, 71^.

BLA input from the lOFC, but not mOFC, was found to mediate reward value encoding. This is consistent with recent evidence that the lOFC itself is necessary for both positive and negative incentive learning ^42^ and with evidence that lOFC lesions disrupt sensitivity of reward seeking to outcome devaluation ^72^. It is also in line with evidence in human lOFC of reward identity coding that is sensitive to reward value shifts ^73, 74^ and identity-based reward value coding ^69^. Interestingly, lOFC→BLA manipulation altered reward value encoding without concomitant changes in the palatability response, consistent with previous evidence of dissociability in these processes ^5^.

That activity at both glutamatergic lOFC terminals and the NMDA receptors known to mediate synaptic plasticity in the BLA is necessary for reward value encoding, suggests that lOFC→BLA projections might direct the encoding of the reward value in the BLA. Consistent with this, an intact lOFC is required for the BLA to encode information about expected outcomes ^75^. Stimulation of lOFC→BLA projections concurrent with reward experience augmented later reward pursuit regardless of deprivation state. Thus, lOFC→BLA projections may convey value to the BLA. Though, this should not rule out a function of the OFC in retaining some forms of reward memory. The lOFC has itself been implicated in the retention and, perhaps, consolidation of action-outcome memories ^76, 77^.

Surprisingly, whereas lOFC→BLA projections were found to mediate reward value encoding, lOFC→BLA activity was neither necessary nor sufficient for retrieving this memory during reward pursuit. Rather, mOFC→BLA projections were found to mediate the retrieval of state-dependent reward value memories. Thus, mOFC→BLA projection activity is critical to ensure reward pursuit commiserate with one’s current state. This is consistent with evidence that the mOFC itself mediates effort allocation according to anticipated reward value ^78^, outcome anticipation ^34^, and other aspects of reward-related decision making ^79, 80^. Confirming that mOFC→BLA projections mediate reward value retrieval, rather than having broader function in reward-related behavior, mOFC→BLA projection manipulation only altered reward pursuit if a state-dependent reward value had been encoded. mOFC→BLA manipulations were without effect in the absence of incentive learning. Moreover, stimulation of mOFC→BLA projections only augmented reward seeking if the internal state was not sufficiently discriminable on its own to support enhanced reward pursuit following incentive learning. Thus, rather than conveying a value signal itself into the BLA, which would have increased reward seeking regardless of prior learning or state, mOFC→BLA projections facilitate the retrieval of reward value, which may be stored in the BLA or downstream.

Both the lOFC and mOFC have been proposed to be involved in representing and using information about the current and anticipated states, or situations, to guide adaptive behavior when the information defining those states (e.g., anticipated reward and its value) is ‘hidden’, i.e., not readily externally observable ^34–36, 81^. For example, the mOFC is necessary for anticipating potential rewarding outcomes and acting accordingly when such outcomes are not present, but is not required not when rewards are themselves present to guide decision making ^34^. Adaptive behavior in our task relies on such a hidden state representation. Although there has been no perceptual change (i.e., same context, levers, etc.), following incentive learning, the state is nonetheless different: the anticipated reward is now more valuable. The critical elements defining this state─ internal need and the reward itself─ are not externally perceptible. Our data, therefore, indicate that lOFC→BLA and mOFC→BLA projections mediate the encoding and retrieval, respectively, of the state-dependent incentive value of a specific anticipated reward.

The demonstrated doubly-dissociable function of lOFC→BLA and mOFC→BLA projections in encoding and retrieving, respectively, of a reward’s value is consistent with recent evidence from primates of similarly dissociable lOFC and mOFC function. The primate lOFC has been shown to be involved in credit assignment ^43–45^ and value updating following devaluation ^46^. Whereas primate mOFC has been implicated in value-guided decision making ^43–45^ and goal selection ^46^. The present results translate this dissociability to rodents and, using bi-directional, projection-specific manipulations, suggest these functions are achieved, at least in part, via projections to the BLA, which are conserved between rat and primate ^22^.

One critical new question is which BLA projections mediate the encoding and retrieval of reward value. Among other potential targets ^3^, the BLA might relay this information back to the OFC. Bilateral BLA lesions will disrupt outcome encoding in the lOFC ^82^, and direct BLA→lOFC projections are necessary for retrieving specific cue-reward memories ^48^. lOFC and mOFC terminals were found to overlap extensively in the BLA, thus, regardless of which BLA neurons mediate reward value memory, the lOFC and mOFC inputs are positioned to target the same network of cells in the BLA.

The contribution of this lOFC→BLA/mOFC→BLA circuitry to other forms of memory is a question ripe for future exploration. Indeed, like the BLA ^4, 83, 3^, the OFC also functions in both appetitive and aversive behavior ^77, 84, 85^ and an intact OFC is necessary for the BLA to encode predicted appetitive or aversive outcomes ^75^. Whether the organizing principle exposed by these data applies to other memory systems is another intriguing possibility. That reward value encoding and retrieval were found to be functionally and neuroanatomically dissociable reveals a clear vulnerability in the brain for poor decision making. Moreover, we found positive reward valuation could be prevented or induced without concomitant changes in the palatability responses indicative of the reward’s emotional experience. OFC-BLA circuitry is known to become dysfunctional in patients diagnosed with addiction ^86, 87^, anxiety ^88, 89^, depression ^90^ and schizophrenia ^91^. The current data, therefore, provide insight into how cortical-amygdala dysfunction might contribute to these and other psychiatric diseases characterized by maladaptive reward valuation and poor reward-related decision making.

## METHODS

### Subjects

Male, Long Evans rats (aged 8–10 weeks at the start of the experiment; Charles River Laboratories, Wilmington, MA) were group housed and handled for 3–5 days prior to the onset of the experiment. Unless otherwise noted, separate groups of naïve rats were used for each experiment. Rats were provided with water *ad libitum* in the home cage and were maintained on food-restriction for a certain amount of time each day, as described below. Experiments were performed during the dark phase of the 12:12 hr reverse dark/light cycle. All procedures were conducted in accordance with the NIH Guide for the Care and Use of Laboratory Animals and were approved by the UCLA Institutional Animal Care and Use Committee.

### Surgery

Standard surgical procedures described previously ^25^ were used for all surgeries. Rats were anesthetized with isoflurane (4–5% induction, 1–2% maintenance) and a nonsteroidal anti-inflammatory agent was administered pre- and post-operatively to minimize pain and discomfort. Following surgery rats were individually housed.

#### Electroenzymatic glutamate recordings

Following training to stable performance, rats were implanted with a unilateral, pre-calibrated glutamate biosensor into the BLA (AP −3.0 mm, ML + 5.1, DV −8.0) and a Ag/AgCl reference electrode into the contralateral cortex. Biosensor placements were verified using standard histological procedures (Fig. 1B).

#### BLA glutamate receptor inactivation

Following training to stable performance, rats were implanted with guide cannula (22-gauge stainless steel; Plastics One, Roanoke, VA) targeted bilaterally 1 mm above the BLA (AP −3.0 mm, ML ± 5.1, DV −7.0). Cannula placements were verified using standard histological procedures (Fig. 1B) and subjects were removed from the study if placements were off target (*N*=1).

#### Chemogenetic manipulation of OFC→BLA projections

Prior to onset of behavioral training, rats were randomly assigned to a OFC subregion group, anesthetized using isoflurane and infused bilaterally with adeno-associated virus (AAV) expressing the inhibitory designer receptor *human M4 muscarinic receptor* (hM4D(Gi); AAV8-CaMKIIa-HA-hM4D(Gi)-IRES-mCitrine). Virus (0.30 µl) was infused at a rate of 6 µl/hr via an infusion needle positioned in the ventrolateral orbitofrontal cortex (lOFC; AP: +3.2 mm; ML: ± 2.4; DV: −5.4) or medial OFC (mOFC; AP: +4.0; ML: ± 0.5; DV: −5.2). Bilateral guide cannula (22-gauge stainless steel; Plastics One) were implanted 1 mm above the BLA (AP −3.0 mm, ML ± 5.1, DV −7.0). Testing commenced 8 weeks post-surgery to ensure axonal transport and expression in lOFC or mOFC terminals in the BLA. Restriction of expression to the lOFC or mOFC was verified with immunofluorescence using an antibody to recognize the HA tag. Cannula placements in the terminal expression region were verified using standard histological procedures. Subjects were removed from the study due to lack of expression or if cannula were misplaced outside the BLA (lOFC, *N*=0, mOFC, *N*=2).

#### Optogenetic manipulation of OFC→BLA projections

Prior to onset of behavioral training, rats were randomly assigned to viral group, anesthetized using isoflurane and infused bilaterally with AAV expressing excitatory opsin channelrhodopsin-2 (ChR2; AAV5-CaMKIIa-hChR2(H134R)-eYFP) or the enhanced yellow fluorescent protein (eYFP) control (AAV8-CaMKIIa-eYFP). Virus (0.30 µl) was infused at a rate of 6 µl/hr via an infusion needle positioned in the lOFC or mOFC. Bilateral optical fibers (200 µm core, numerical aperture 0.66; Prizmatix, Southfield, MI) held in ferrules (Kientec Systems Inc., Stuart, FL) were implanted 0.3 mm above the BLA (AP −3.0 mm, ML ± 5.1, DV −7.7). Testing commenced 8 weeks post-surgery to ensure axonal transport and expression in lOFC or mOFC terminals in the BLA. Restriction of virus to either the lOFC or mOFC was verified with immunofluorescence using an antibody against eYFP and optical fiber placements in vicinity of terminal expression were verified using standard histological procedures. Subjects were removed from the study due to lack of expression or if optical fibers were misplaced outside the BLA (lOFC, *N*=1, mOFC, *N*=1).

#### Validation of chemogenetic and optogenetic manipulation of OFC→BLA projections

hM4d(Gi) and ChR2 were co-expressed by infusing both AAV5-CaMKIIa-hChR2(H134R)-eYFP and either AAV8-CaMKIIa-HA-hM4D(Gi)-IRES-mCitrine or AAV5-CaMKIIa-mCherry bilaterally in the lOFC (AP: +3.2 mm; ML: ± 2.4; DV: −5.4) or mOFC (AP: +4.0; ML: ± 0.5; DV: −5.2). 8-weeks post viral infusion, rats were anesthetized and a pre-calibrated microelectrode array (MEA) glutamate biosensor affixed to an optical fiber and guide cannula was acutely implanted into the BLA (AP −3.0 mm, ML + 5.1, DV −8.0), and a Ag/AgCl reference electrode placed in the contralateral cortex. The optical fiber was affixed behind the MEA (to reduce photovoltaic artifacts) and the optical fiber tip terminated 0.3 mm above the glutamate sensing electrodes. The guide cannula (Plastics One) terminated 6.5 mm above the MEA tip to avoid tissue damage and was positioned such that, when inserted, the injector (Plastics One) would protrude 6.2 mm and end within 100 µm from the microelectrodes. The injector was inserted after the biosensor/optical fiber probe was lowered into the BLA to further minimize tissue damage. Level of anesthesia was kept constant throughout recordings by maintaining breaths per minute (bpm) constant (1 bpm) by adjusting isoflurane level (1–1.5%). Viral expression was verified using immunofluorescence and biosensor placements were verified using standard histological procedures (Fig. S10).

### Electroenzymatic glutamate biosensors

#### Biosensor fabrication

MEA probes were fabricated in the Nanoelectronics Research Facility at UCLA and modified for glutamate detection as described previously ^23–25^. Briefly, these biosensors use glutamate oxidase (GluOx) as the biological recognition element and rely on electro-oxidation, via constant-potential amperometry (0.7 V versus a Ag/AgCl reference electrode), of enzymatically-generated hydrogen peroxide reporter molecule to provide a current signal. This current output is recorded and converted to glutamate concentration using a calibration factor determined *in vitro*. Enzyme immobilization was accomplished by chemical crosslinking using a solution consisting of GluOx, bovine serum albumin (BSA), and glutaraldehyde. Interference from both electroactive anions and cations is effectively excluded from the amperometric recordings, while still maintaining a subsecond response time, by electropolymerization of polypyrrole (PPY) or poly o-phenylenediamine (PPD), as well as dip-coat application of Nafion to the electrode sites prior to enzyme immobilization ^23–25^. Each MEA had two non-enzyme-coated sentinel electrodes for the removal of correlated noise from the glutamate sensing electrodes by signal subtraction, as described previously ^24, 25^. These electrodes were prepared identically with the exception that the BSA/glutaraldehyde solution did not contain GluOx. The average *in vivo* limit of glutamate detection of the sensors used in this study was 0.36 µM (sem=0.03, range 0.13–0.67 µM).

#### Reagents

Nafion (5 wt.% solution in lower aliphatic alcohols/H_2_O mix), bovine serum albumin (BSA, min 96%), glutaraldehyde (25% in water), pyrrole (98%), p-Phenylenediammine (98%), L-glutamic acid, L-ascorbic acid, 3-hydroxytyramine (dopamine) were purchased from Aldrich Chemical Co. (Milwaukee, WI, USA). L-Glutamate oxidase (GluOx) from *Streptomyces* Sp. X119–6, with a rated activity of 24.9 units per mg protein (U mg^−1^, Lowry’s method), produced by Yamasa Corporation (Chiba, Japan), was purchased from US Biological (Massachusetts MA). Phosphate buffered saline (PBS) was composed of 50 mM Na_2_HPO_4_ with 100 mM NaCl (pH 7.4). Ultrapure water generated using a Millipore Milli-Q Water System (resistivity = 18 MΩ cm) was used for preparation of all solutions used in this work.

#### Instrumentation

Electrochemical preparation of the sensors was performed using a Versatile Multichannel Potentiostat (model VMP3) equipped with the ‘p’ low current option and low current N’ stat box (Bio-Logic USA, LLC, Knoxville, TN). *In vitro* and *in vivo* measurements were conducted using a low-noise multichannel Fast-16 mkIII potentiostat (Quanteon LLC, Nicholasville, KY), with reference electrodes consisting of a glass-enclosed Ag/AgCl wire in 3 M NaCl solution (Bioanalytical Systems, Inc., West Lafayette, IN) or a 200 µm diameter Ag/AgCl wire, respectively. All potentials are reported versus the Ag/AgCl reference electrode. Oxidative current was recorded at 80 kHz and averaged over 0.25-s intervals.

#### In Vitro Biosensor Characterization

All biosensors were calibrated *in vitro* to test for sensitivity and selectivity of glutamate measurement prior to implantation. A constant potential of 0.7 V was applied to the working electrodes against a Ag/AgCl reference electrode in 40 mL of stirred PBS at pH 7.4 and 37ºC within a Faraday cage. After the current detected at the electrodes equilibrated (~30–45 min), aliquots of glutamate were added to the beaker to reach final glutamate concentrations in the range 5 – 60 µM. A calibration factor based on these response was calculated for each GluOx-coated electrode. The average calibration factor for the sensors used in these studies was 135.98 µM/nA. Control electrodes, coated with PPy or PPD, Nafion, and BSA/glutaraldehyde, but not GluOx, showed no detectable response to glutamate. Aliquots of ascorbic acid (250 µM final concentration) and dopamine (5–10 µM final concentration) were added to the beaker as representative examples of readily oxidizable potential anionic and cationic interferent neurochemicals, respectively, to confirm selectivity for glutamate (Fig. S1). For the sensors used in these studies no current changes above the level of the noise were detected to the addition of cationic or anionic interferents, as reported previously ^23–25^. To assess uniformity of H_2_O_2_ sensitivity across control and GluOx-coated electrodes, aliquots of H_2_O_2_ (10 µM) were also added to the beaker. There was less than a 10%, statistically insignificant (t_42_=0.32, p=0.75) difference in the H_2_O_2_ sensitivity on control electrode sites relative to enzyme-coated sites, indicating that any changes detected *in vivo* on the enzyme-coated biosensor sites following control channel signal subtraction could not be attributed to endogenous H_2_O_2_.

### In vivo validation of chemogenetic and optogenetic manipulation of OFC→BLA projections

Glutamate biosensors were used to validate optogenetic stimulation and chemogenetic inhibition, respectively, of OFC terminals in the BLA. Animals expressing ChR2 and hM4d(Gi) in either the lOFC or mOFC were anesthetized and implanted with a pre-calibrated MEA-fiber-cannula probe into the BLA, as described above. Experiments were conducted inside a Faraday cage. Following sensor implant, an injector was inserted into the cannula. A constant potential of 0.7 V was applied to the working electrodes against the Ag/AgCl reference electrode implanted in the contralateral hemisphere. The detected current was allowed to equilibrate (~30–45 min). Baseline spontaneous glutamate release events (i.e., glutamate transients) were measured for 2 min prior to infusion of vehicle. Spontaneous transients were then monitored for 15 min post-infusion. Following this, glutamate release was optically evoked by delivery of blue light pulses (473 nm, 5–20 mW, 20Hz Hz, 5 s or 3 s) to stimulate lOFC or mOFC terminals in the BLA. Each stimulation parameter was repeated 3x, with at least 60 s in between stimulations. Rats then received an infusion of CNO (1 mM, 0.5 µl) into the extracellular space surrounding the MEA. Spontaneous glutamate transients were monitored 2 min before (baseline) and 15 min following CNO infusion. The light delivery protocol was then repeated to assess the effect of CNO:hM4D(Gi) or CNO:mCherry of optically-evoked glutamate release from OFC terminals in the BLA. As an iterative control, in a subset of subjects, the applied potential was lowered to 0. 2 V, below the H_2_O_2_ oxidizing potential, and recordings of spontaneous and optically-evoked glutamate release were made following CNO infusion.

### Optical stimulation

Light was delivered to the OFC terminals in the BLA using a laser (Dragon Lasers, ChangChun, JiLin, China) connected through a ceramic mating sleeve (Thorlabs, Newton, NJ) to the ferrule implanted on the rat. We used a 473 nm laser to activate ChR2-transfected projection neurons, or a 589 nm laser (which is largely outside the range of the ChR2 sensitivity range ^92^) as a control for the effects of construct expression and light delivery in ChR2-transfected projection neurons. For optical stimulation, light pulses (25 msec pulse) were delivered at 20 Hz. This was based on previous studies showing reward-induced firing rates of OFC neurons that range from 6–40 spikes/second ^84, 93–95^. We also found this stimulation frequency to effectively stimulate glutamate release from OFC terminals in the BLA *in vivo* (Fig. S10). Light effects were estimated to be restricted to the BLA based on predicted irradiance values (https://web.stanford.edu/group/dlab/cgi-bin/graph/chart.php).

### Drug Administration

Ifenprodil (Tocris Bioscience, Bristol, UK) and NBQX (2,3-Dioxo-6-nitro-1,2,3,4-tetrahydrobenzo[*f*]quinoxaline-7-sulfonamide disodium salt; Tocris Bioscience, Bristol, UK) were dissolved in sterile saline vehicle. CNO (Tocris Bioscience, Bristol, UK) was dissolved in aCSF to 1mM. Drugs were infused bilaterally into the BLA in a volume of 0.5 µl over 1 min via injectors inserted into the guide cannula fabricated to protrude 1 mm ventral to the cannula tip using a microinfusion pump. Injectors were left in place for at least 1 additional min to ensure full infusion. This infusion volume was selected to avoid spread to the adjacent central nucleus of the amygdala ^5^. Rats were placed in the conditioning chamber 5 min after infusion to allow sufficient time for the drug to become effective. The dose of ifenprodil (1.67 µg/side), an N-methyl-D-aspartate (NMDA) receptor antagonist with selective targeting of receptors that contain the NR2B subunit ^30^, was selected because it has been shown to impair value-based decision making ^7^. The alpha-amino-3-hydroxyl-5-methyl-4-isoxazole-propionate (AMPA) receptor antagonist, NBQX, at a dose of 1.0 µg/side, was selected based on our previous evidence of its effectiveness in reward-related tasks ^25, 96^. CNO dose was selected based on our previous demonstration of the efficacy and duration of action of this dose and our evidence showing effective inhibition of glutamate release from OFC terminals in the BLA with this dose (Fig. S10) ^48^. We have also demonstrated that this dose of CNO when infused into the BLA has no effect on reward-related behavior in the absence of the hM4D(Gi) transgene ^48^.

### Behavioral Procedures

#### General training and testing

##### Apparatus

Training took place in Med Associates conditioning chambers (East Fairfield, VT) housed within sound- and light-attenuating boxes, described previously ^25^. For *in vivo* glutamate measurements all testing was conducted in a single Med Associates conditioning chamber housed within a continuously-connected, copper mesh-lined sound attenuating chamber and outfitted with an electrical swivel (Crist Instrument Co, Hagerstown, MD) connecting a headstage tether that extended within the conditioning chamber to the potentiostat recording unit (Fast-16 mkIII, Quanteon, LLC) positioned outside the conditioning chamber. For optogenetic experiments, testing was conducted in Med associates conditioning chambers outfitted with an Intensity Division Fiberoptic Rotary Joint (Doric Lenses, Quebec, QC, Canada) connecting the output fiberoptic patchcords to a laser (Dragon Lasers, ChangChun, JiLin, China) positioned outside the conditioning chamber.

All chambers contained 2 retractable levers that could be inserted to the left and right of a recessed food-delivery port in the front wall. A photobeam entry detector was positioned at the entry to the food port to provide a goal approach measure. The chambers were equipped with syringe pump to deliver 20% sucrose solution in 0.1ml increments through a stainless steel tube, or a pellet dispenser that delivered a single 45-mg pellet (Bio-Serv, Frenchtown, NJ), into a custom-designed electrically-isolated Acetal plastic well in the food port. A lickometer circuit (Med Associates), connecting the grid floor of the boxes and stainless steel sucrose-delivery tubes, with the circuit closed by the rats’ tongue allowed recording of the lick frequency (licks/s) when rats consumed each sucrose delivery. A 3-watt, 24-volt house light mounted on the top of the back wall opposite the food-delivery port provided illumination.

##### Training

Each experiment followed the same general structure. Rats were trained on a self-paced, 2-lever, action sequence to earn a delivery of 0.1ml 20% sucrose. Training procedures were similar to those we have described previously ^5, 6, 26^. Except where noted, rats were deprived of food for 4 hr prior to each training session. Each session began with the illumination of the houselight and insertion of the lever, where appropriate, and ended with the retraction of the lever and turning off of the houselight. Rats were given only one training session/day. Rats received 3 d of magazine training in which they were exposed to non-contingent sucrose or water deliveries (30 outcomes over 35 min) in the conditioning chamber with the levers retracted, to learn where to receive sucrose. This was followed by daily instrumental training sessions in which sucrose could be earned by lever pressing. Rats were first given 3 d of single-action, taking lever, instrumental training on the lever to the right (i.e., ‘taking’ lever) of the food-delivery port with the sucrose delivered on a continuous reinforcement schedule. Each session lasted until 20 outcomes had been earned or 30 min elapsed. Following single-action instrumental training, the ‘seeking’ lever (i.e., the lever to the left of the food-delivery port) was introduced into the chamber. Rats were allowed to press on the seeking lever to gain access to the taking lever, a single press on which delivered the sucrose solution and retracted this lever. The seeking lever remained present during the entire session. Rats were trained on this self-paced, 2-lever, action sequence for a total of 12–18 days: 3 days in which a press on the ‘seeking’ lever was continuously reinforced with the taking lever, 2–4 days in which the seeking lever was reinforced on a random ratio 2 (RR-2) schedule, 3–5 days in which the seeking lever was reinforced on a RR-5 schedule, and 4–6 days in which the seeking lever was reinforced on the final RR-10 schedule until stable responding was established. The taking lever was always continuously reinforced. Each session lasted until 20 outcomes had been earned or 40 min elapsed.

##### Incentive learning opportunity and test

Following training to stable response rates, rats received non-contingent re-exposure to the sucrose outcome (30 exposures/35 min) in the conditioning chamber with the levers retracted. Unless otherwise noted, food-port entries and lickometer palatability measures ^97–100^ were collected during this phase of the experiment. These non-contingent sucrose deliveries provided an incentive learning opportunity wherein the value of the sucrose may be updated (see specific experimental procedures). The sucrose re-exposure was non-contingent to avoid any caching of value to the action itself. The next day lever-press behavior was measured during a brief, 5-min, non-reinforced probe test to assess the effects of the previous day’s incentive learning opportunity on reward-seeking actions. Because no sucrose was delivered during this test, there was no opportunity for online incentive learning or new reinforcement learning. Thus, this task allowed us to experimentally isolate reward value encoding v. reward value retrieval.

##### Online, near-real time glutamate detection during sucrose exposure or seeking

Following training on the self-paced action sequence in the sated (4-hr food-deprived) state and surgery (see Fig. 1a), testing commenced. Prior to each test, rats were placed in the recording conditioning chamber and the biosensor was tethered to the potentiostat via the electrical swivel for application of the 0.7 V potential. The recorded amperometric signal was allowed to stabilize prior to session onset (~30–45 min). First, rats received a single day of instrumental re-training similar to the training described above, but with the ratio requirement progressively increasing from a fixed-ratio-1 to RR-10 after each 5^th^ outcome earned to re-establish lever pressing post-surgery. The next day, rats were non-contingently exposed to the sucrose in the familiar sated state (4 hr food-deprived) or in a hungry (20 hr food-deprived) state. For hunger state group assignment, subjects were counterbalanced based on average lever-press rate during the last 2 instrumental training sessions. The next day, all rats were tested hungry. A separate group of rats were maintained hungry throughout training and test (Fig. S5). To prevent electrical interference, lickometers were not connected during recording sessions.

##### BLA AMPA and NMDA glutamate receptor inactivation during sucrose re-exposure or post-re-exposure lever-pressing test

Following training in the sated state as described above, drug groups were counterbalanced based on lever-press rate during the two final instrumental training sessions. On 2 of the instrumental training days immediately prior to the first incentive learning opportunity rats were given mock infusions to habituate them to the infusion procedures; injectors were inserted into the cannula, but no fluid was infused. All rats then received the non-contingent re-exposure to the sucrose in the 20 hr food-deprived hungry state. Prior to this incentive learning opportunity, rats received intra-BLA infusions of vehicle, Ifenprodil, or NBQX. The next day all rats received a drug-free, non-reinforced, lever-pressing probe test in the hungry state (see Fig. 2a). Following 2 days to reestablish satiety, rats received two sessions of retraining (1/day) on the action sequence in the 4-hr food-deprived state. They were then given another round of re-exposure and a lever-pressing test. In this case, non-contingent exposure to the sucrose in the hungry state was conducted drug-free. To ensure value encoding and to equate the number of incentive learning opportunities with intact glutamate receptor activity, rats previously assigned to the vehicle group received 2 drug-free re-exposure sessions, while the rats previously assigned to Ifenprodil or NBQX groups received 3 drug-free re-exposure sessions. The day following the last day of re-exposure, all rats received a non-reinforced, lever-pressing probe test in the hungry state. Prior to this test, rats received an infusion of vehicle, Ifenprodil, or NBQX (see Fig. 2f). Rats received the same drug on both tests.

##### Chemogenetic inactivation of lOFC→BLA or mOFC→BLA projections during sucrose re-exposure or post-re-exposure lever-pressing test

Training and test was identical to that for the BLA glutamate receptor inactivation experiments, except that rats expressing hM4D(Gi) in the lOFC or mOFC received infusion of either vehicle or CNO. All rats received mock infusions to habituate them to the infusion procedures. Following training, rats received non-contingent re-exposure to the sucrose in the 20 hr food-deprived hungry state. Prior to this incentive learning opportunity, rats received intra-BLA infusions of either vehicle or CNO. The next day all rats received a drug-free, non-reinforced, lever-pressing probe test in the hungry state (see Fig. 3a). Following 2 days to reestablish satiety, rats received two sessions of retraining (1/day) on the action sequence in the 4-hr food-deprived state. They were then given another round of re-exposure. In this case, non-contingent exposure to the sucrose in the hungry state was conducted drug-free. Rats previously assigned to the vehicle group received 2 drug-free re-exposure sessions, while the rats previously assigned to the CNO groups received 3 drug-free re-exposure sessions. The day following the last day of re-exposure, all rats received a non-reinforced, lever-pressing probe test in the hungry state immediately following infusion of either vehicle or CNO into the BLA (see Fig. 2G). Drug group assignment for this test was counterbalanced with respect to previous drug treatment. There was no effect of previous drug group (*F*_1,24_=1.51, *P*=0.23) or interaction between this variable and experimental group (*F*_2,24_=0.93, *P*=0.41) on reward-seeking during this test, indicating that the results of this test were not influenced by drug history. There were no significant differences in reward-seeking lever presses between vehicle-treated subjects expressing hM4D(Gi) in the lOFC or mOFC during either the first (*t_11_*=2.00, p=0.07) or second test (*t_9_*=0.20, p=0.85), and therefore, these groups were collapsed to serve as a single control group.

To evaluate the effect of mOFC→BLA projection inactivation on reward seeking in the absence of reward value retrieval, a separate group of rats expressing hM4D(Gi) in the mOFC was trained sated and received intra-BLA infusions of vehicle or CNO prior to a non-reinforced, lever-pressing probe test in the hungry state as above, but without prior non-contingent re-exposure to the sucrose in the hungry state (i.e., without a reward value encoding opportunity; Fig. S13). Each rat was given 2 non-reinforced probe tests, one each following vehicle or CNO infusion for a within-subject drug comparison (test order counterbalanced). Two days after the last non-reinforced probe test, rats were retrained sated for two days, given a drug-free incentive learning opportunity in the hungry state, and then received intra-BLA infusions of Vehicle or CNO prior to a reinforced lever-pressing test (Fig. S13). In this test, the presence of the sucrose made retrieval of its value from memory unnecessary. Each rat was given 2 reinforced tests, one each following vehicle or CNO infusion to allow a within-subject drug comparison (test order counterbalanced).

##### Optogenetic activation of OFC→BLA projections during sucrose re-exposure

Rats expressing ChR2, or the eYFP control in the lOFC or mOFC with optical fibers above the BLA were trained sated as described above (Fig. 4a). On the last two days of instrumental training, rats were tethered to the patchcord, but no light was delivered to allow habituation to the optical tether. At test, rats were maintained in the familiar 4-hr food-deprived sated state and received non-contingent re-exposure to the sucrose or to a task-irrelevant food-pellet. During this non-contingent sucrose exposure blue light (473 nm, 20Hz, 10mW, 5 s) was delivered for optical activation of lOFC terminals, in the BLA in ChR2-expressing subjects, during consumption of the sucrose. The laser was triggered by the first lick following sucrose delivery or the first food-port entry following pellet delivery. Optical stimulation timing was based on evidence that BLA glutamate release occurred in response to sucrose consumption during incentive learning and peaked on average 2.79 s (s.e.m.=0.67; range = 0.63–6.1 s) post sucrose collection (Fig. 1d) and evidence that rats finish sucrose consumption and exited the food delivery port ~5–10 s following reward collection. A subset of rats expressing ChR2 received 589 nm light delivery (outside the range of ChR2 sensitivity ^92^) in the BLA. The next day, all rats received a non-reinforced probe test in the familiar sated state while tethered, but without light delivery. This sequence of re-exposure and testing was repeated twice, first in a novel, moderate hunger state (8-hr food-deprived) and then in a novel hungry (20-hr food-deprived) state (Fig. S15). Rats were given 2 days off and retrained in the 4-hr food-deprived state for two days in between each test set. In no case did reward-seeking lever-press activity significantly differ between ChR2-expressing rats that received 589 nm optical activation and eYFP controls receiving 473 nm optical activation (*t^6^*=0.10–0.95, p=0.38–0.93) and, thus, these controls groups were collapsed to serve as a single control group for each test.

##### Optogenetic activation of OFC→BLA projections during lever-pressing test

Rats expressing ChR2, or eYFP in the lOFC or mOFC with optical fibers above the BLA received training, non-contingent sucrose exposure, and testing as the described above, except light (473 nm, 20Hz, 10mW, 3 s) was delivered during each of the non-reinforced lever-pressing tests to, in ChR2-expressing subjects, activate lOFC or mOFC terminals in the BLA. Light was delivered 1/minute, for a total of 10 light deliveries throughout the 10-minute test. The first light delivery occurred 30 s after test onset. The duration of optical stimulation was based on the finding that glutamate release preceded the initiation of reward seeking and the rise time to peak glutamate release prior to reward-seeking bouts was on average 1.95 s (s.e.m.=0.43; range = 0.40–3.0 s; Fig. 1f). As above, a subset of ChR2-expressing subjects received 589 nm light delivery. Tests were conducted 4-, 8-, and 20-hr food-deprived, as above, with each pressing test preceded by non-contingent sucrose re-exposure in the absence of light delivery. The moderate 8-hr food-deprived state provided a subthreshold incentive learning opportunity that was, on its own, not sufficiently discriminable to induce an upshift in reward seeking. Reward-seeking presses did not significantly differ between ChR2-expressing rats that received 589 nm light delivery and eYFP controls receiving 473 nm light delivery (*t_6_*=0.30–2.44, p=0.051–0.77) and, thus, these groups were collapsed to serve as a single control group for each test.

To examine the effect of mOFC→BLA projection activation on reward seeking in the moderate food-deprivation state, but in the absence of incentive learning, a separate group of rats expressing ChR2 in the mOFC was trained while sated, and received light delivery during a non-reinforced probe test in the moderate 8-hr food-deprived state as above, but without prior re-exposure to the sucrose in the 8-hr food-deprived state (i.e., without the subthreshold incentive learning opportunity). Each rat was given 2 non-reinforced probe tests, one each with either 473 nm (for ChR2 activation) or 589 nm (control wavelength) light delivery, to allow within-subject comparison. Test order was counterbalanced across subjects.

### Histology

Rats were transcardially perfused at the conclusion of behavioral testing with PBS followed by 10% formalin. The brains were removed, post-fixed in formalin, then cryoprotected, cut with a cryostat at a thickness of 30 µm, and collected in PBS. eYFP fluorescence was used to verify ChR2 expression. To verify hM4D(Gi) expression, immunohistochemical analysis was performed as described previously ^101–103^. Briefly, floating coronal sections were blocked for 1 hr at room temperature in 8% normal goat serum (NGS, Jackson ImmunoResearch Laboratories) with 0.3% Triton X-100 in PBS and then incubated overnight at 4°C in 2% NGS, 0.3% Triton X-100 in PBS with primary antibody (anti-HA, 1:500, Biolegend, San Diego, CA, cat. no. 901501). The sections were then incubated for 2 hr at room temperature with goat anti-mouse IgG, Alexa 594 conjugate (1:1000, Invitrogen, cat. no. A11005). All sections were washed 3 times for 5 min each in PBS before and after each incubation step and mounted on slides using ProLong Gold antifade reagent with DAPI (Invitrogen). All images were acquired using a Keyence (BZ-X710) microscope with a 4X or 20X objective (CFI Plan Apo), CCD camera, and BZ-X Analyze software. Biosensor and cannula placements in non-AAV subjects, were verified using standard histological procedures.

### Data analysis

#### Behavioral analysis

Seeking and taking lever presses and/or food-port entries collected continuously for each training and test session. Seeking lever presses were normalized to baseline response rate averaged across the last 2 training sessions prior to test to control for pre-test response variability and allow comparison across tests conducted in different deprivation states (see ^5, 6, 104, 105^). Raw press rates data are presented in the supplemental materials. Lickometer measurements were made during sucrose consumption during the non-contingent re-exposure sessions.

#### Chemogenetic and optogenetic manipulation of glutamate release

Analysis details and characterization of glutamate release events have been described previously ^24, 25^. Electrochemical data were baseline-subtracted. Detected current was averaged across the first 10 s of the 2-min, pre-infusion, baseline period and this baseline was subtracted from current output at each time point. Current changes from baseline on the PPY(or PPD)/Nafion-coated sentinel electrode were then subtracted from current changes on the PPY(or PPD)/Nafion/GluOx glutamate biosensor electrode to remove correlated noise. This signal was then converted to glutamate concentration using an electrode-specific calibration factor obtained *in vitro*. Mini Analysis (Synaptosoft, Decatur, GA) was used to determine the frequency and amplitude of spontaneous glutamate transient release events. A fluctuation in the glutamate trace was deemed a glutamate transient if it was at >2.5x the RMS noise sampled from the pre-test baseline period. To determine transient amplitude, a baseline was taken by averaging 3 sample bins around the first minima located 0.5–5 s before the peak and this baseline was subtracted from the peak amplitude. If one peak followed another within 5 s the baseline was taken after the first peak to distinguish these events. Peaks with a total duration below 0.5 s or with an immediately preceding or following negative deflection greater than half the peak amplitude were considered noise spikes and were omitted from the analysis. To evaluate optically-evoked glutamate release, we isolated the 5-s or 3-s period prior to, during, and following light delivery. The average glutamate concentration change in the 5-s or 3-s optical stimulation period was subtracted from that during an equivalent period immediately prior to optical stimulation. This was averaged across each of the 3 replicates for each parameter. There were no statistically-significant main effects of OFC subregion (mOFC v. lOFC; *F*_1,4_=2.09, *P*=0.22; Treatment: *F*_1,4_=8.78, *P*=0.04; Brain region × Treatment: *F*_1,4_=0.01, *P*=0.91)) and, thus, these data were collapsed.

#### Temporal relationship between glutamate release and behavior

As above, electrochemical data were baseline-subtracted. Detected current was averaged across the 10 s baseline period 2-min prior to test and this baseline was subtracted from current output at each time point. We evaluated the temporal relationship between glutamate release and behavioral events as described previously ^24, 25^. For the sucrose re-exposure, we isolated glutamate concentration changes in the 5 s prior to and 10 s following the first food-port entry following each sucrose delivery (i.e., reward collection). This period was chosen to give an adequate pre-sucrose baseline and based on evidence that rats disengaged from the food port ~5–10 s following sucrose collection. The average glutamate concentration in the 1-s period 5 s prior to sucrose collection served as the baseline and this was subtracted from each data point in the peri-sucrose glutamate concentration v. time trace. To quantify the sucrose-evoked glutamate concentration change, for each trial the average glutamate concentration change in the 10-s post-sucrose period was averaged across trials and this was compared to average glutamate concentration change in the 5-s prior to sucrose collection and to equivalent analysis of glutamate concentration changes in 5-s periods in the absence of sucrose or checking behavior.

During the non-reinforced, lever-pressing probe test, because rats tended to organize their reward-seeking lever presses into bouts, we focused on those presses that initiated bouts of reward-seeking activity (i.e., ‘initiating presses’), excluding presses that occurred within a pressing bout, as we have described previously ^25^. An ‘initiating seeking press’ was defined as the first press after completion of an action-sequence or, because rats often disengaged from the lever and then reinitiated reward seeking, the first press after ≥6 s pause in pressing. Similar definitions of initiation of reward seeking and instrumental bouts defined by pauses in activity have been described previously ^25, 106–108^. See Table S1 for seeking bout information. We evaluated glutamate concentration changes in the 5 s prior to and following each initiating reward-seeking press. The average glutamate concentration in the 1-s period, 5 s prior to each initiating press served as the baseline. This analysis window was selected to avoid contaminating events (e.g., termination of a previous bout, food-port entries, etc.). Average glutamate concentration change for each initiating press was quantified in the 3-s period immediately prior to and after each initiating press and this was compared to equivalent analysis of glutamate concentration changes in the absence of lever pressing. Data were averaged across trials. We quantified glutamate concentration around all intra-bout seeking presses similarly (Fig. S6). Pearson correlations were used to assess the relationship between glutamate fluctuations around bout initiation and the number of presses and duration of subsequent bouts.

#### Palatability analysis

A lickometer circuit (Med Associates), connecting the grid floor of the box and the stainless steel sucrose-delivery tubes, with the circuit closed by the rats’ tongue, allowed recording of individual lick events. Lickometer measures were amplified and fed through an interface to a PC programmed to record the time of each lick to the nearest 1 msec. Based on previous reports ^5, 97, 104, 109–111^, we used licking frequency (licks/sec) as a measure of sucrose palatability. This measure of licking microstructure during consumption provides a similar analysis of palatability changes as those assessing taste reactivity following oral infusions ^99^. These data were analyzed with custom-written python-based code.

#### Statistical analysis

Datasets were analyzed by two-tailed, Student’s *t* tests, one- or two-way repeated-measures analysis of variance (ANOVA), as appropriate. Bonferroni corrected *post hoc* tests were performed to clarify all main effects and interactions. Two-tailed, paired t-tests were used for *a priori* planned comparisons, as advised by ^112^ based on a logical extension of Fisher’s protected least significant difference (PLSD) procedure for controlling familywise Type I error rates. All datasets met equal covariance assumptions, justifying ANOVA interpretation ^113^. Alpha levels were set at *P*<0.05.

## DATA AVAILABILITY

All data that support the findings of this study are available from the corresponding author.

## ACKNOWLEDGEMENTS

This research was supported by NIH grant DA035443, MH106972, and NS087494 to KMW and NIH grant DA038942 and DA024635 to MM. We would like to acknowledge the helpful feedback from Nina Lichtenberg and Dr. Alicia Izquierdo on these data and this manuscript.

## AUTHOR CONTRIBUTIONS

MM and KMW designed the research, analyzed, and interpreted the data. MM conducted the research with assistance from CS, MDM, and VYG. MM and KMW wrote the manuscript.

## Supplemental Figures

**Table S1.** Summary of seeking lever press bouts during non-reinforced, lever-pressing probe test. Data presented as average ± s.e.m.

| Exposure Group | Number of bouts<br>(per 5-min test) | Average<br>presses per bout | Average<br>bout duration (s) |
| --- | --- | --- | --- |
| 4-hr | 9.33±1.26 | 2.35±0.33 | 3.24±0.76 |
| 20-hr | 9.67±1.69 | 5.86±2.35 | 6.10±2.41 |

**Supplement Figure 1.**
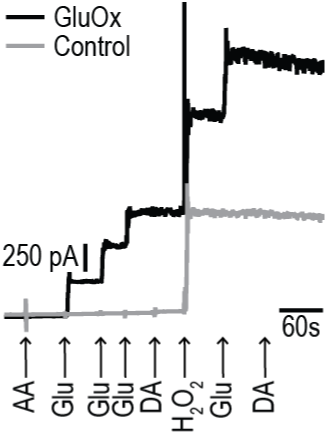
Representative calibration of a microelectrode array glutamate biosensor. Silicon-wafer-based platinum microelectrode array (MEA) probes were modified for glutamate detection as we have described previously ^24, 25, 114^. Glutamate oxidase (GluOx) serves as the biological recognition. Electro-oxidation, by constant potential amperommetry, of the enzymatically-generated hydrogen peroxide reporter molecule provides the signal. Selectivity against both cations and anions is achieved by the addition of polymer coatings (see Methods). Control electrodes are identically coated with the exception that GluOx is omitted. These sensors have a subsecond response time ^24, 25, 114^. To test for sensitivity and selectivity of glutamate measurement, all biosensors were calibrated *in vitro* by sequential addition of ascorbic acid (AA; 250μM), glutamate (Glu; 20 μM), dopamine (DA; 5 μM), hydrogen peroxide (H_2_O_2_; 20 μM), Glu (40 μM), and DA (10 μM), in stirred PBS at 37 ^o^C. The *in vitro* glutamate current response was used to determine the electrode-specific calibration factor, which averaged 135.98 µM/nA for the sensors used in these studies. The sensitivity to peroxide between glutamate oxidase coated (GluOx) and control sites did not differ more than 10% (t_42_=0.32, p=0.75). The average *in vivo* limit of glutamate detection of the sensors used in this study was 0.36 µM (sem=0.03, range 0.13–0.67 µM).

**Supplement Figure 2.**
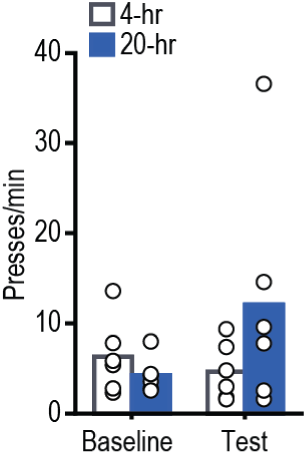
Effect of incentive learning on reward seeking-raw press rates. Reward-seeking press rate (seeking presses/min) during baseline (average of last-two training sessions in 4-hr food-deprived state prior to test) and non-reinforced, lever-pressing probe test in the hungry state (Test: *F*_1,10_=3.1.577, *P*=0.24; Deprivation: *F*_1,10_=0.71, *P*=0.42; Test × Deprivation: *F*_1,10_=3.73, *P*=0.08) for rats prior non-contingent sucrose exposure in control sated (4-hr food-deprived; no value encoding) or hungry (20-hr deprived; value encoding opportunity) state. Data presented as mean + scatter.

**Supplement Figure 3.**
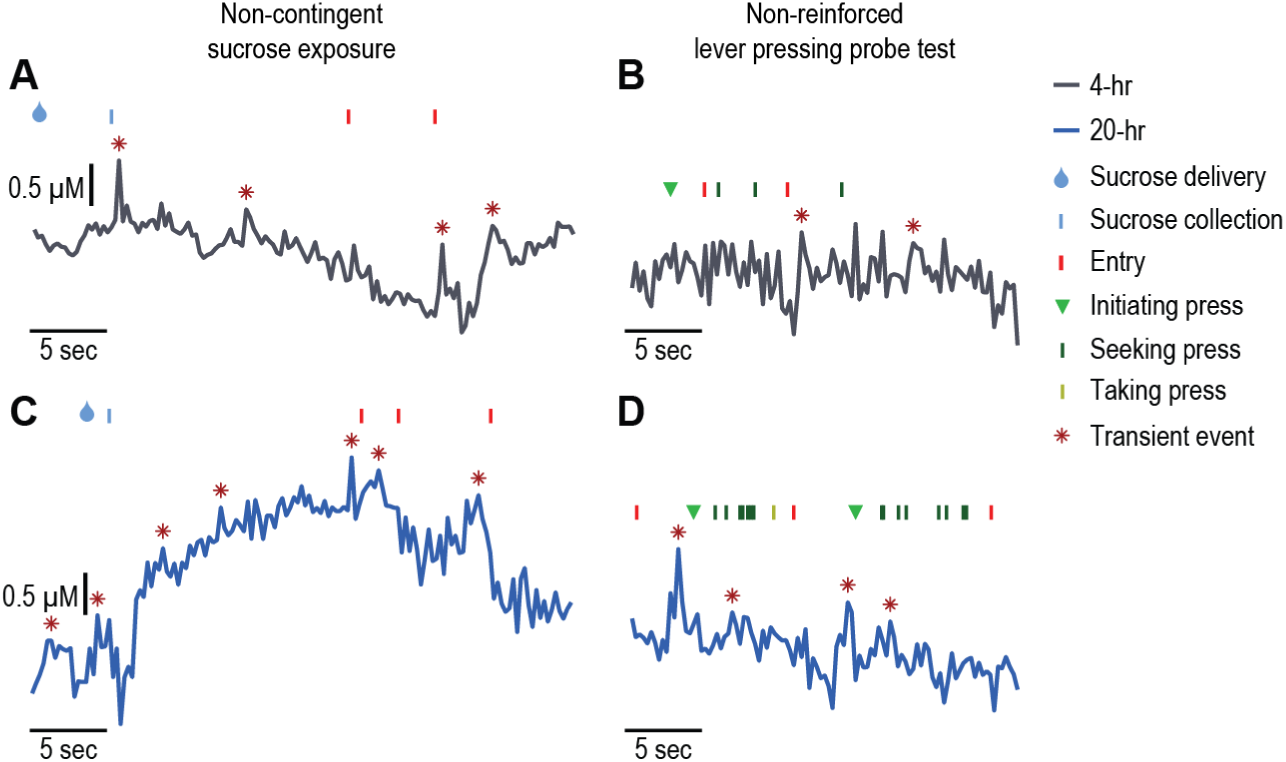
Representative BLA glutamate v. time traces during reward value encoding and retrieval. Representative, single-trial BLA glutamate concentration v. time traces from a rat that received non-contingent sucrose re-exposure in (**a-b**) the control, familiar sated (4-hr), or (**c-d**) the novel hungry (20-hr; positive value encoding opportunity) state around (**a, c**) sucrose collection during the non-contingent re-exposure and (**b, d**) the subsequent lever-pressing activity in the non-reinforced probe test conducted in the hungry state.

**Supplement Figure 4.**
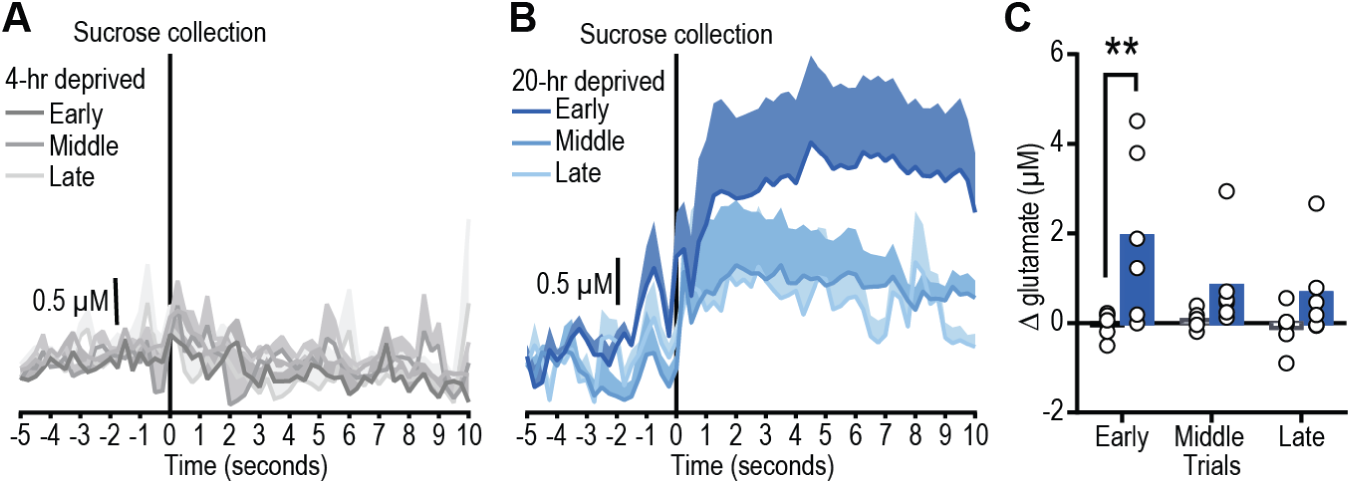
BLA glutamate release during non-contingent reward re-exposure - binned. (**a-b**) Trial-averaged, BLA glutamate v. time trace (shading reflects between-subject s.e.m) around sucrose collection and (**c**) quantification (mean + scatter) of average glutamate immediately post reward consumption during early (1–10), middle (11–20), or late (21–30) reward-delivery trials (**a**) in the familiar sated (4-hr food deprived) state or (**b**) novel hungry (20-hr food-deprived, incentive learning opportunity) state (Trial bin: *F*_2,20_=3.70, *P*=0.04; Deprivation: *F*_1,10_=5.52, *P*=0.04; Trial bin × Deprivation: *F*_2,20_=3.81, *P*=0.04). Sucrose-evoked glutamate release is largest early in the re-exposure session, when incentive learning is expected to be the highest.

**Supplement Figure 5.**
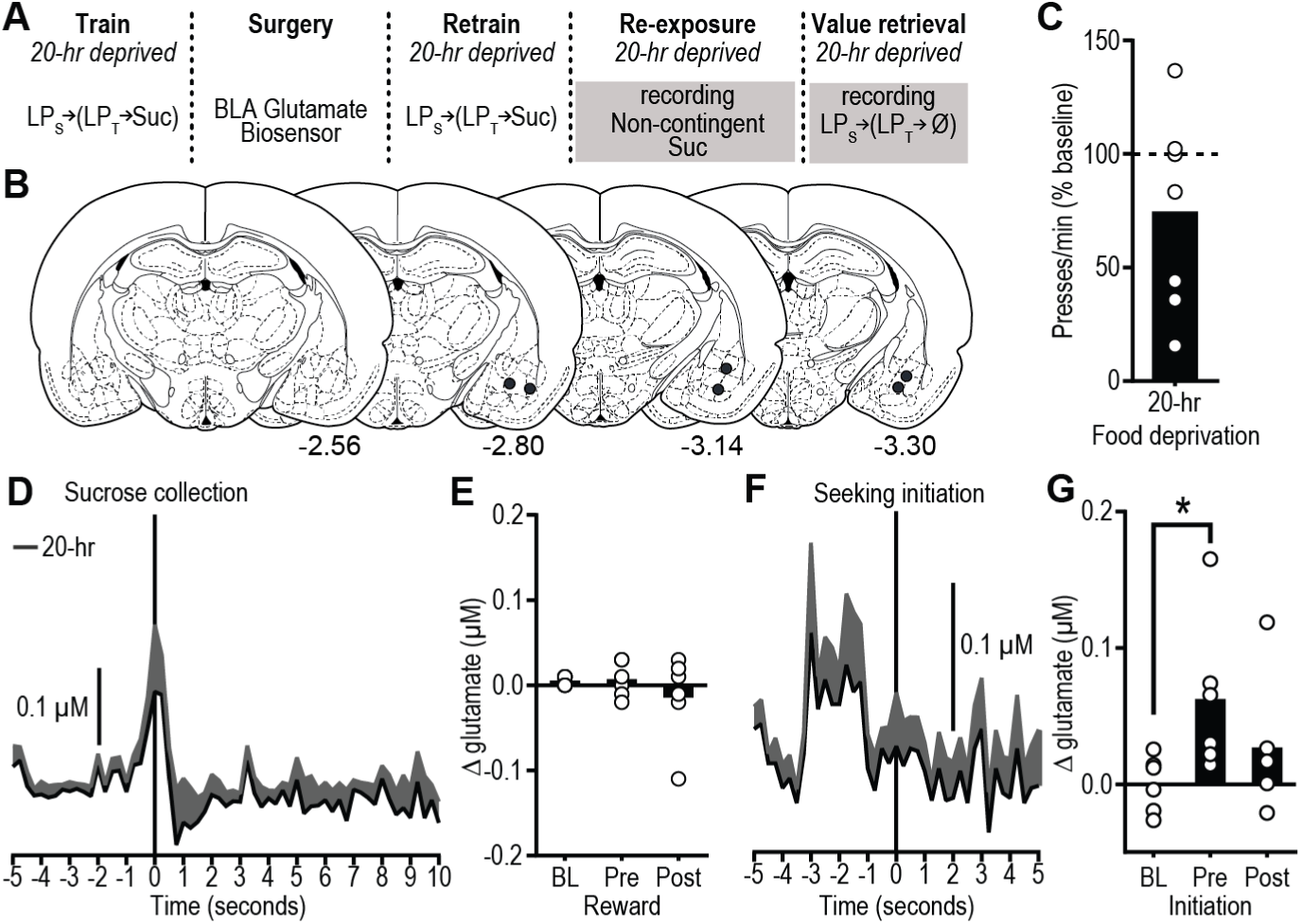
BLA glutamate release during reward value encoding and retrieval in familiar hungry state. (**a**) Procedure schematic (LPs, seeking lever press; LPt, taking lever press; Suc, sucrose; Ø, no sucrose delivered). Rats were trained while hungry (20-hr food-deprived) to press on the seeking-taking chain to earn sucrose. Biosensor glutamate recordings were made during non-contingent re-exposure to the sucrose in the familiar hungry state and during a lever-pressing probe test, also in the hungry state. (**b**) Placement of the microelectrode array biosensor tips in BLA. Numbers represent anterior-posterior distance (mm) from bregma. (**c**) Reward-seeking press rate (seeking presses/min), relative to baseline press rate (dashed line), during non-reinforced, lever-pressing probe test in the hungry (20-hr food-deprived) state. (**d**) Trial-averaged BLA glutamate concentration v. time trace (shading reflects between-subject s.e.m.) and (**e**) quantification (mean + scatter) of average glutamate concentration change prior to (pre) and following (post) sucrose collection/consumption (occurring at time 0 s), or equivalent baseline periods (BL) during non-contingent sucrose re-exposure in familiar hungry state (*N*=6; *F*_2,10_=0.86, *P*=0.409). (**f**) Trial-averaged BLA glutamate concentration v. time trace and (**g**) quantification of average glutamate concentration change around bout-initiating reward-seeking presses during the lever-pressing probe test in the hungry state (*F*_2,10_=4.13, *P*=0.049). \**p* < 0.05, relative to baseline. Reward experience in the hungry state does not increase BLA glutamate concentration in the absence of encoding new information about the value of the reward in that state.

**Supplement Figure 6.**
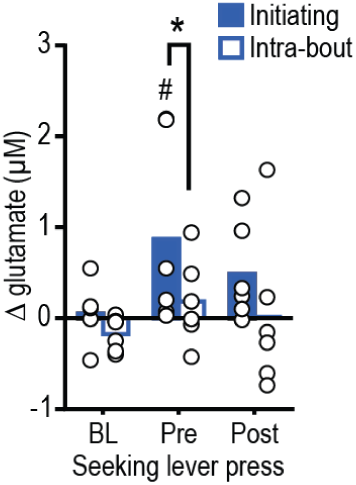
BLA glutamate concentration around all reward-seeking presses. Quantification of average glutamate concentration change around all seeking presses, separating intra-bout presses from bout-initiating seeking presses, during the lever-pressing probe test in the hungry state for subjects that had prior incentive learning experience with the sucrose in the hungry state (Time: *F*_2,10_=3.07, *P*=0.09; Press type: *F*_1,5_=8.15, *P*=0.04; Time × Press type: *F*_2,10_=0.96, *P*=0.42). \**P*<0.05, between groups; ^#^*P*<0.05, relative to baseline. Data presented as mean + scatter. Glutamate transients do not precede each individual press but rather only precede bout-initiating reward-seeking presses.

**Supplement Figure 7.**
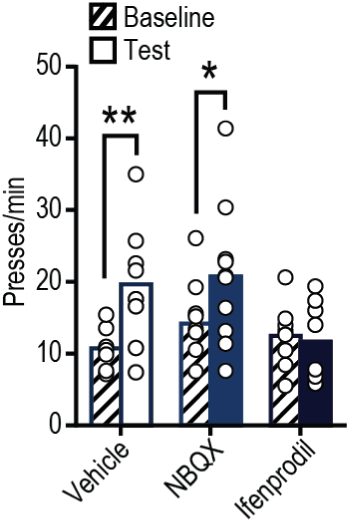
Effect of glutamate receptor antagonist on value encoding - raw press rates. Reward-seeking press rate (seeking presses/min) during baseline and drug-free non-reinforced lever-pressing probe test in the hungry state (Test: *F*_1,23_=12.57, *P*=0.002; Treatment: *F*_2,23_=2.01, *P*=0.16; Test × Treatment: *F*_2,23_=4.31, *P*=0.03) in rats that received BLA microinfusion of vehicle, AMPA, or NMDA antagonist during prior non-contingent sucrose exposure in hungry (20-hr) state. \**P*<0.05, \*\**P*<0.01 compared to baseline. Data presented as mean + scatter.

**Supplement Figure 8.**
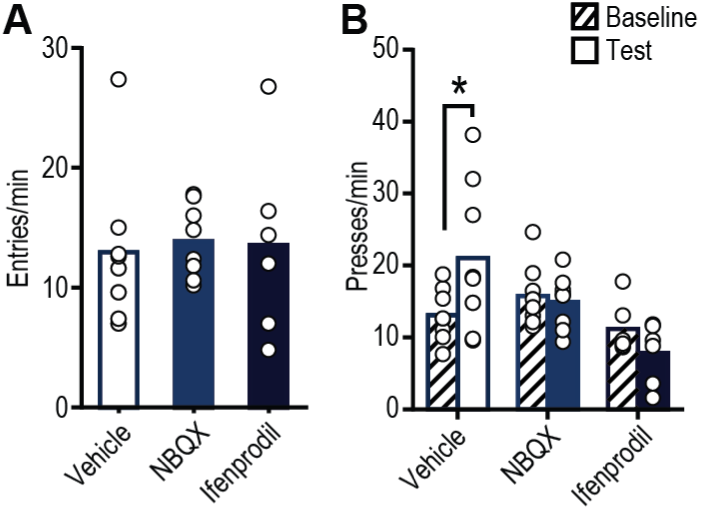
Effect of glutamate receptor antagonists on reward checking and reward seeking - raw entry/press rates. Following non-contingent sucrose exposure in hungry (20-hr food-deprived) state, rats received intra-BLA of Vehicle, AMPA, or NMDA antagonist prior to a non-reinforced, lever-pressing probe test in the hungry state. (**a**) Food-port entry rate (entries/min; *F*_2,19_=0.06, *P*=0.95) during this test. Neither treatment affected this reward-checking measure. (**b**) Reward-seeking press rate (seeking presses/min) during baseline and the on-drug post-re-exposure, non-reinforced, lever-pressing probe test (Test: *F*_1,19_=0.69, *P*=0.42; Treatment: *F*_2,19_=4.95, *P*=0.02; Test × Treatment: *F*_2,19_=5.44, *P*=0.01). *\*P<*0.05, relative to baseline. Data presented as mean + scatter.

**Supplement Figure 9.**
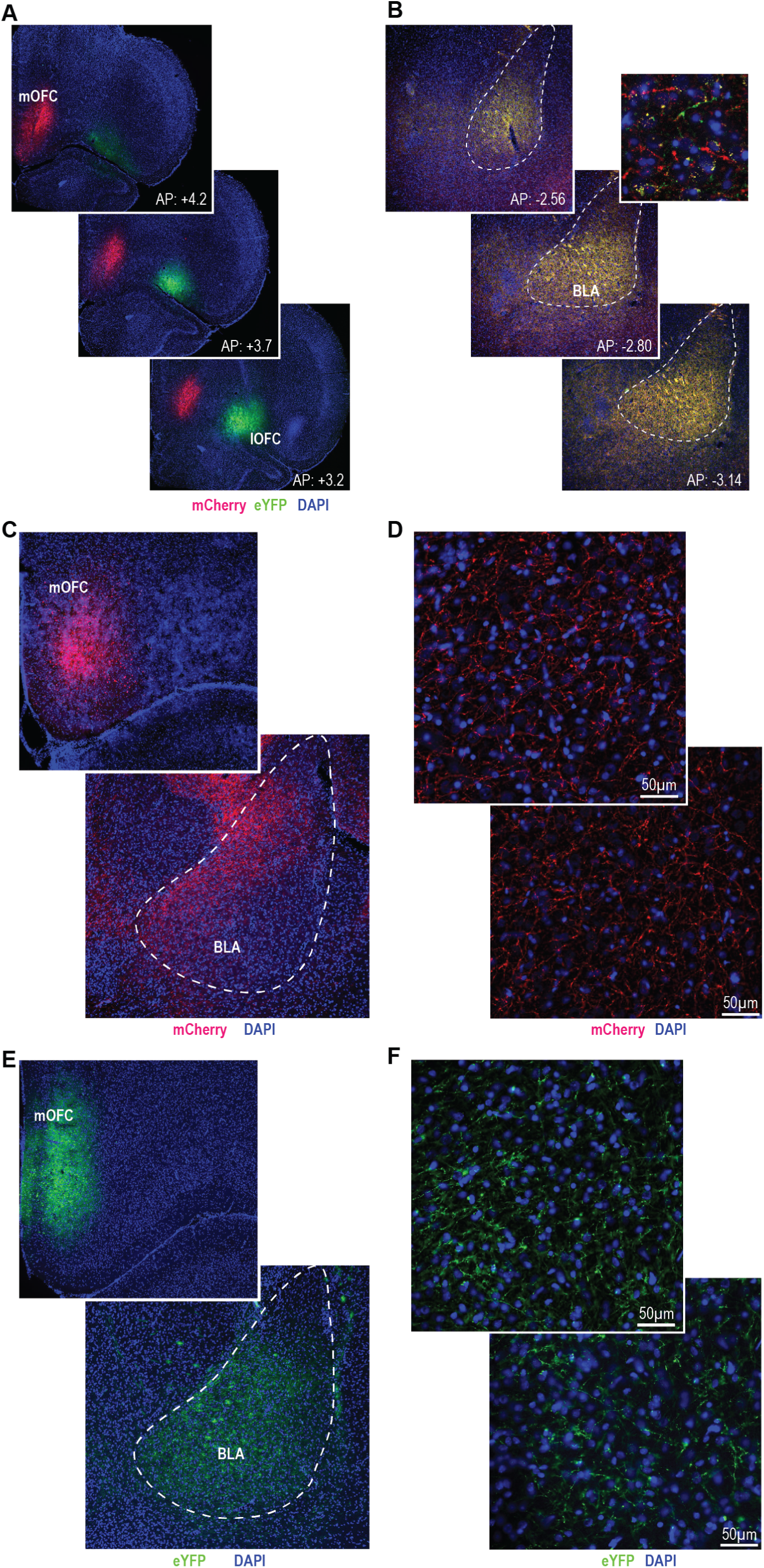
lOFC and mOFC projections to the BLA. (**a-b**) AAV5-CaMKIIa-mCherry was infused in the mOFC and AAV5-CaMKIIa-eYFP was infused into the lOFC and allowed to express for 8 weeks, to ensure terminal expression, prior to histological assessment. (**a**) Representative expression of mCherry and eYFP in the mOFC and lOFC, respectively. (**b**) Expression of mCherry and eYFP restricted to fibers in the BLA. These data provide anatomical evidence of intermingled projections from both the ventrolateral OFC and medial OFC to the BLA. (**c-f**) AAV5-CamKIIa-ChR2-eYFP or AAV8-hSyn-hM4D(Gi)-mCherry was infused into the mOFC and allowed to express for 8 weeks. (**c, e**) Representative expression of eYFP (**c**) or mCherry (**e**) in the mOFC infusion location and the BLA terminal field. (**d, f**) Expression of eYFP (**e**) or mCherry (**f**) restricted to fibers in the BLA.

**Supplement Figure 10.**
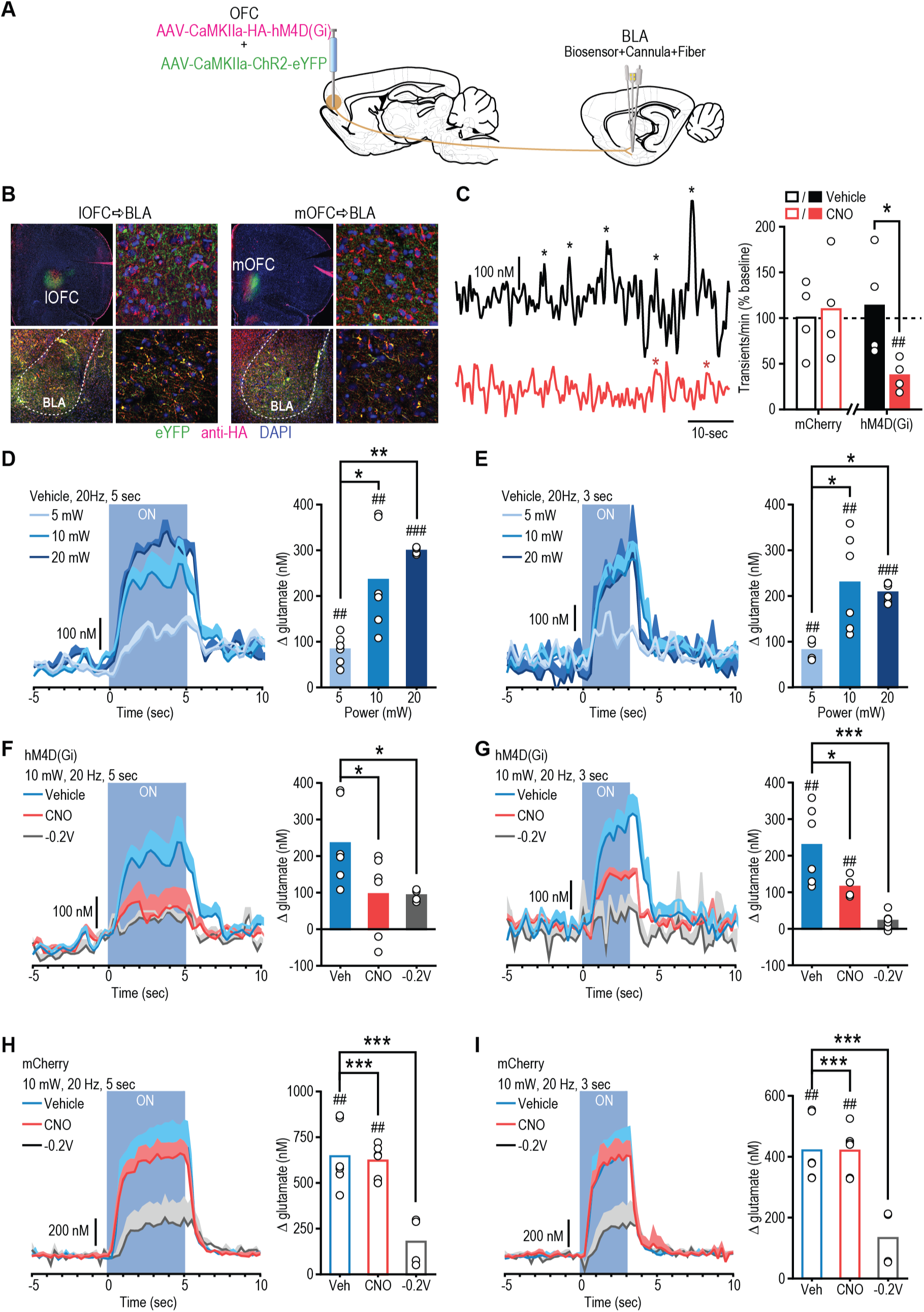
Chemogenetic and optogenetic manipulation of OFC terminals in BLA. (**a**) Procedure schematic. hM4d(Gi) and ChR2 were co-expressed in either the lOFC or mOFC. Following 8 weeks for terminal expression, we measured spontaneous and optically-evoked glutamate release events in the BLA of anesthetized rats prior to and following CNO infusion. (**b**) Representative immunofluorescent images of HA-tagged hM4D(Gi) and eYFP-tagged ChR2 expression in lOFC (*left*) or mOFC (*right*) and BLA terminal expression (*bottom*). Data from lOFC and mOFC subjects was collapsed following evidence of no statistically significant differences between these groups. (**c**) Representative glutamate concentration v. time trace showing spontaneous, transient glutamate release events following vehicle or CNO (1mM/0.5µl) treatment and quantification of glutamate transient rate (transients/min) normalized to pre-infusion baseline rate (dashed line) (*t*^6^=2.54, *p*=0.04). These data indicate that chemogenetic inhibition of OFC terminals can decrease spontaneous glutamate release events in the BLA. Importantly, however, this should not be taken as evidence that the OFC contributes to ~50% of spontaneous BLA activity because such activity is dependent on a variety of BLA inputs and interneurons that are likely differentially sensitive to anesthesia and the animal’s current state. (**d-e**) Optically-evoked BLA glutamate concentration v. time trace (shading reflects s.e.m.) and quantification (mean + scatter) of optically-evoked glutamate concentration changes. Blue light delivery for (**d**) 5-sec (*F*_2,13_=11.65, *P*=0.001) or (**e**) 3-sec (*F*_2,13_=6.34, *P*=0.01) over OFC terminals in the BLA power-dependently evoked a glutamate concentration change. (**f-g**) Glutamate concentration v. time trace around (**f**) 5-sec (*F*_2,13_=3.77, *P*=0.05) or (**g**) 3-sec (*F*_2,13_=13.80, *P*>0.001) optical stimulation of OFC terminals in BLA following intra-BLA Vehicle or CNO infusion and quantification. Optically-evoked response following CNO did not differ from current changes detected below the H_2_O_2_ (glutamate reporter molecule) oxidizing potential (0.2 V). (**h-i**) In a separate group of subjects, ChR2 was co-expressed with mCherry to control for non-specific effects of CNO in the absence of the hM4D(Gi) transgene. Glutamate concentration v. time trace around (**h**) 5-sec (*F*_2,13_=14.91, *P*>0.001) or (**i**) 3-sec (*F*_2,13_=16.16, *P*>0.001) optical stimulation of OFC terminals in BLA following intra-BLA Vehicle or CNO infusion and quantification. Optically-evoked response following CNO did not differ from current changes detected following vehicle infusion in subjects lacking hM4D(Gi). \**P*<0.05, ** *P*<0.01, *** *P*<0.001, between groups; ^#^*P*<0.05, ^##^*P*<0.01 relative to baseline. See also ^48^ for additional validation of OFC→BLA chemogenetic and optogenetic terminal manipulations with *ex vivo* electrophysiology.

**Supplement Figure 11.**
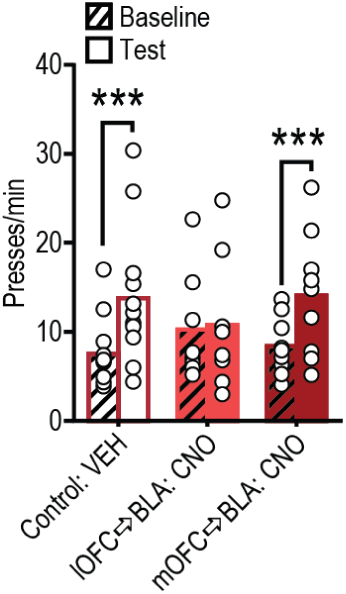
Effect of inactivation of lOFC or mOFC terminals in the BLA on reward value encoding - raw press rates. Reward-seeking press rate (seeking presses/min) during baseline and drug-free, non-reinforced lever-pressing probe test in the hungry state (Test: *F*_1,26_=22.94, *P*<0.0001; Treatment: *F*_2,26_=0.04, *P*=0.96; Test × Treatment: *F*_2,26_=4.21, *P*=0.03) for rats that received BLA microinfusion of Vehicle or CNO during the non-contingent sucrose re-exposure in the hungry (20-hr food-deprived) state. \*\**P*<0.01, \*\*\**P*<0.001, relative to baseline. Data presented as mean + scatter.

**Supplement Figure 12.**
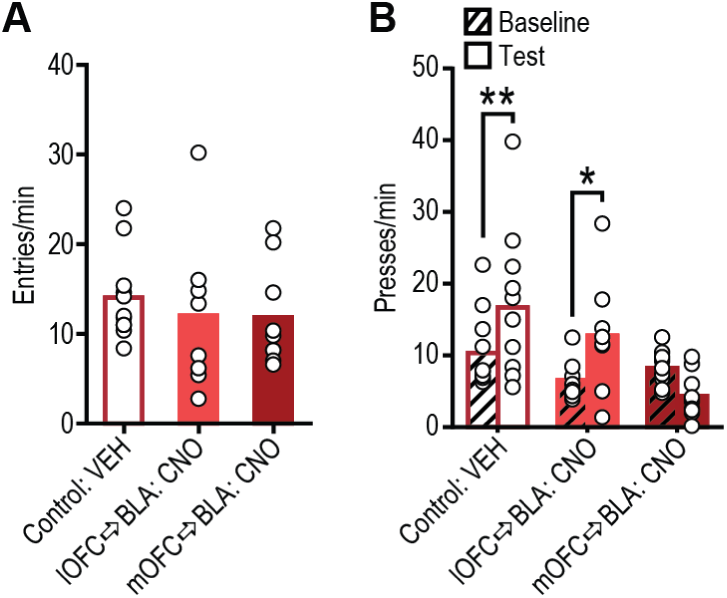
Effect of inactivation of lOFC or mOFC terminals in the BLA on reward-checking and reward seeking - raw entry/press rates. Following non-contingent sucrose exposure in hungry (20-hr food-deprived) state, rats received BLA microinfusion of vehicle or CNO prior to a non-reinforced lever-pressing probe test in the hungry state. (**a**) Food-port entry rate (entries/min) was not altered by inactivation of either lOFC or mOFC terminals in the BLA during this test (*F*_2,25_=0.36, *P*=0.70). (**b**) Reward-seeking press rate (seeking presses/min) during baseline and the on-drug post-re-exposure, non-reinforced, lever-pressing probe test (Test: *F*_1,25_=6.54, *P*=0.02; Treatment: *F*_2,25_=4.30, *P*=0.02; Test × Treatment: *F*_2,25_=8.94, *P*=0.001). \**P*<0.05, \*\**P*<0.01, relative to baseline. Data presented as mean + scatter.

**Supplement Figure 13.**
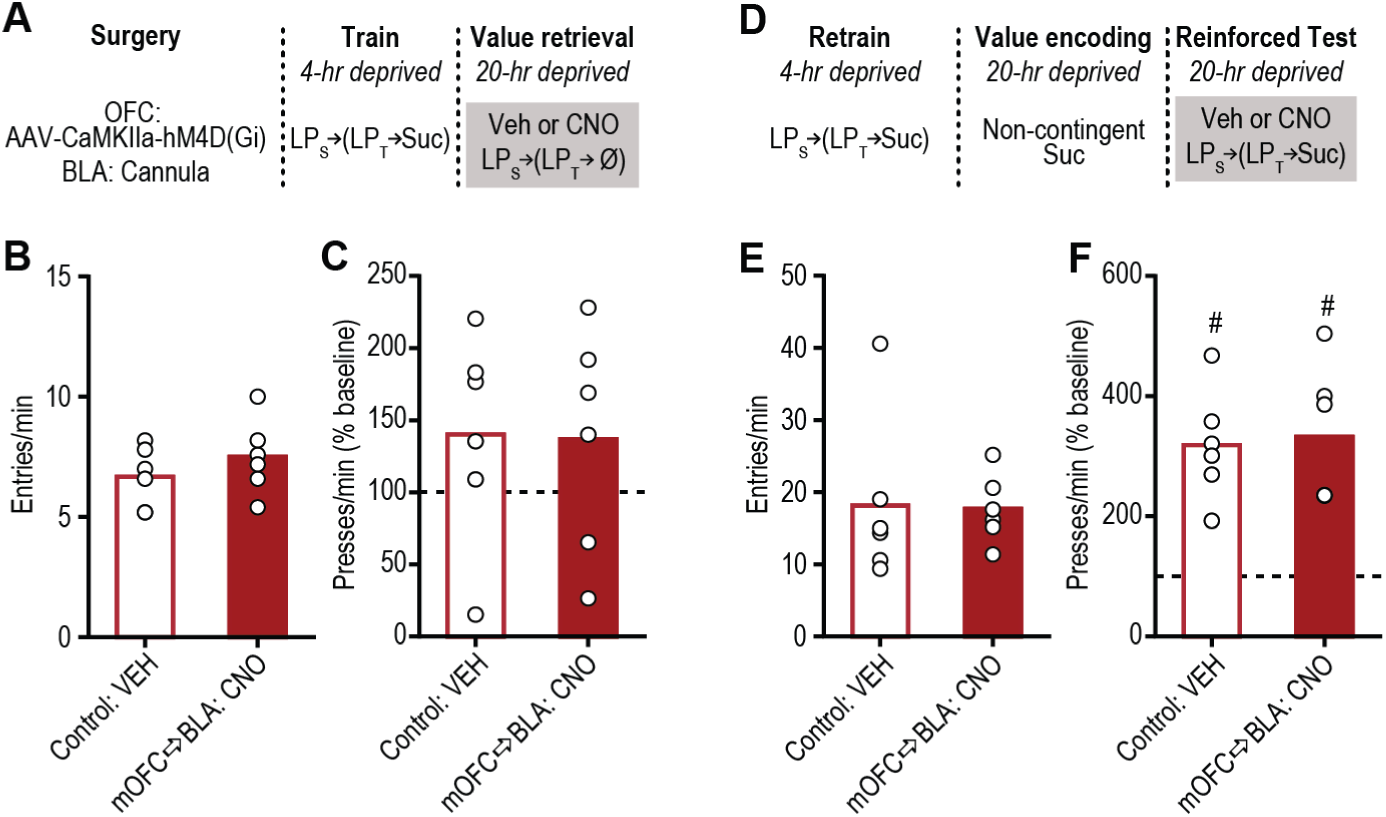
Inactivation of mOFC terminals in the BLA does not disrupt reward seeking when reward value is not being retrieved from memory. (**a**) Procedure schematic. Rats were trained while sated to lever press on the seeking-taking chain to earn sucrose. Following training, they were given two non-reinforced, lever-pressing, probe tests in the hungry state- one each following intra-BLA vehicle or CNO infusion. (LPs, seeking lever press; LPt, taking lever press; Suc, sucrose; Ø, no sucrose delivered; Veh, vehicle; CNO; Clozapine N-oxide) (**b**) Food-port entry rate (entries/min) (*t*_5_=1.01, *p*=0.36) and (**c**) reward-seeking press rate (seeking presses/min), normalized to baseline press rate (dashed line), during the non-reinforced lever-pressing probe test in the hungry state following BLA microinfusion of Vehicle or CNO (*N*=6). mOFC→BLA terminal inactivation was ineffective at altering reward-seeking activity in the absence of prior hunger-induced incentive learning (*t*_5_=0.09, *p*=0.93). (**d**) Procedure schematic. Following retraining in the sated state, rats were given non-contingent re-exposure to the sucrose in the hungry state (the incentive learning opportunity) and then were given two reinforced lever-pressing tests in the hungry state, one each following BLA vehicle or CNO infusion (order counterbalanced). (**e**) Food-port entry rate (entries/min) (*t*_5_=0.15, *p*=0.89) and (**f**) reward-seeking press rate (seeking presses/min), relative to baseline press rate (dashed line). (*N*=6) mOFC→BLA terminal inactivation was ineffective at altering reward-seeking activity if reward value had been encoded, but did not have to be retrieved because the reward was present at test (*t*_5_=0.34, *p*=0.75). ^#^*P*<0.05, relative to baseline. Data presented as mean + scatter.

**Supplement Figure 14.**
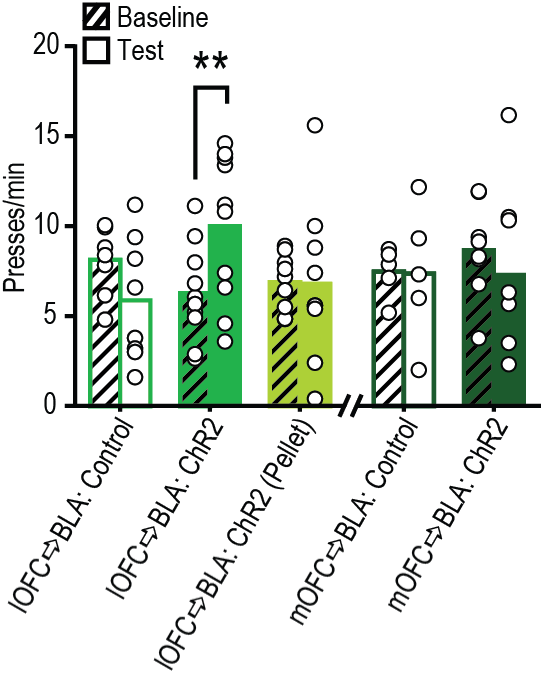
Effect of optical stimulation of lOFC→BLA or mOFC→BLA projections on value encoding - raw press rates. Reward-seeking press rate (seeking presses/min) during baseline and the post-re-exposure, manipulation-free, non-reinforced, lever-pressing probe test in the sated state in rats that received prior non-contingent sucrose or task-irrelevant (Pellet) exposure concurrent with light delivery during in sated state. lOFC→BLA: Test: *F*_1,24_=0.54, *P*=0.47; Group: *F*_2,24_=0.60, *P*=0.55; Test × Group: *F*_2,24_=7.89, *P*=0.002; mOFC→BLA: Test: *F*_1,11_=0.49, *P*=0.50; Group: *F*_1,11_=0.11, *P*=0.74; Test × Group: *F*_1,11_=0.36, *P*=0.56. \*\**P*<0.01, relative to baseline. Data presented as mean + scatter.

**Supplement Figure 15.**
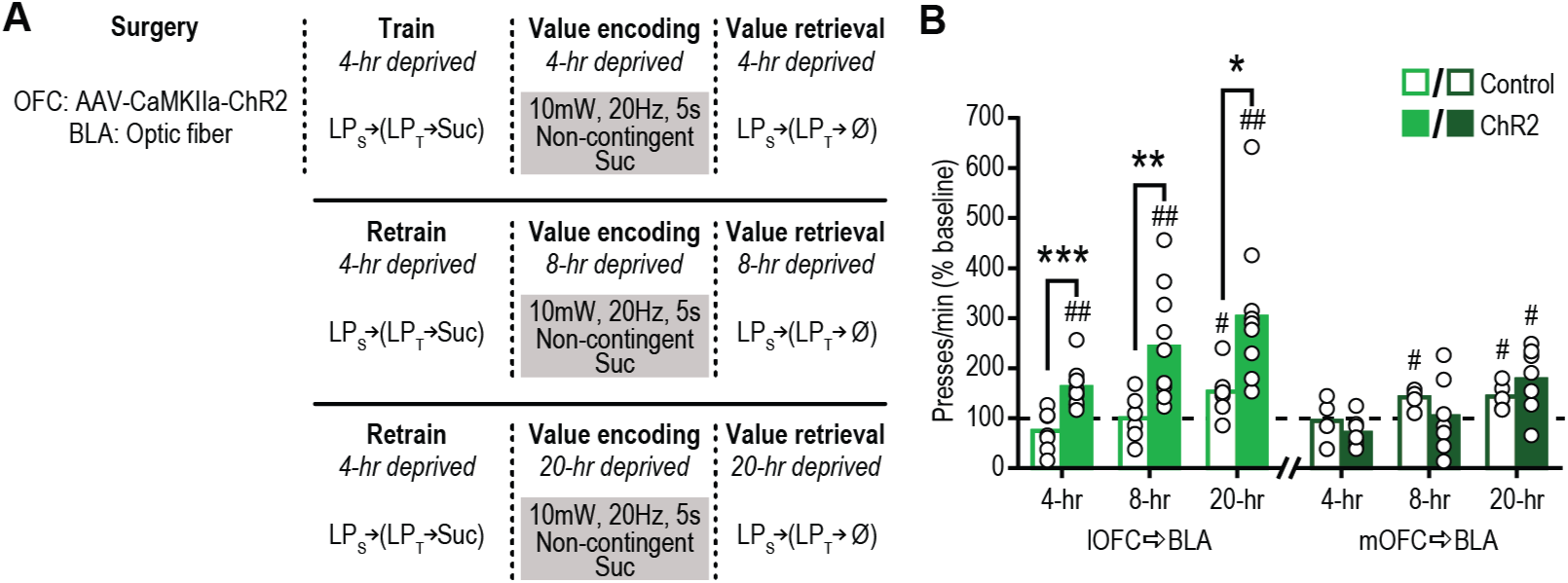
Activation of lOFC, but not mOFC, to BLA projections concurrent with sucrose experience is sufficient to enhance value assignment across escalating food-deprivation states. (**a**) Procedure schematic. Rats received 3 test sets in which they first received non-contingent re-exposure to the sucrose with concurrent optical activation of lOFC or mOFC terminals in BLA (ChR2 + 473 nm, 10mW, 20Hz, 5 s) or control light delivery (control group consisted of half eYFP + 473 nm and half ChR2 + 589 nm light delivery), and then, the next day, received a non-reinforced, lever-pressing probe test in the same deprivation state. Rats were tested at escalating food-deprivation levels (control, familiar 4-hr food deprived state, moderate 8-hr food-deprived state, and hungry 20-hr food-deprived state). (**b**) Reward-seeking press rate (seeking presses/min), relative to baseline press rate (dashed line), during the non-reinforced, lever-pressing probe test conducted the day following non-contingent sucrose re-exposure. At each deprivation state tested, activation of lOFC terminals in the BLA concurrent with non-contingent sucrose-experience caused a subsequent upshift in reward-seeking activity (Group: *F*_1,15_=20.74, *P*=0.0004; deprivation: *F*_2,30_=7.46, *P*=0.002; Group × deprivation: *F*_2,30_=0.73, *P*=0.49). Activation of mOFC terminals in the BLA concurrent with non-contingent sucrose-experience did not alter subsequent reward seeking, compared to controls (Group: *F*_1,10_=0.32, *P*=0.59; deprivation: *F*_2,20_=6.61, *P*=0.006; Group × deprivation: *F*_2,20_=1.62, *P*=0.22). Planned comparisons: \**P*<0.05, \*\**P*<0.01, \*\*\**P*<0.001, between groups; ^#^*P*<0.05 ^##^*P*<0.01, relative to baseline. Data presented as mean + scatter.

**Supplement Figure 16.**
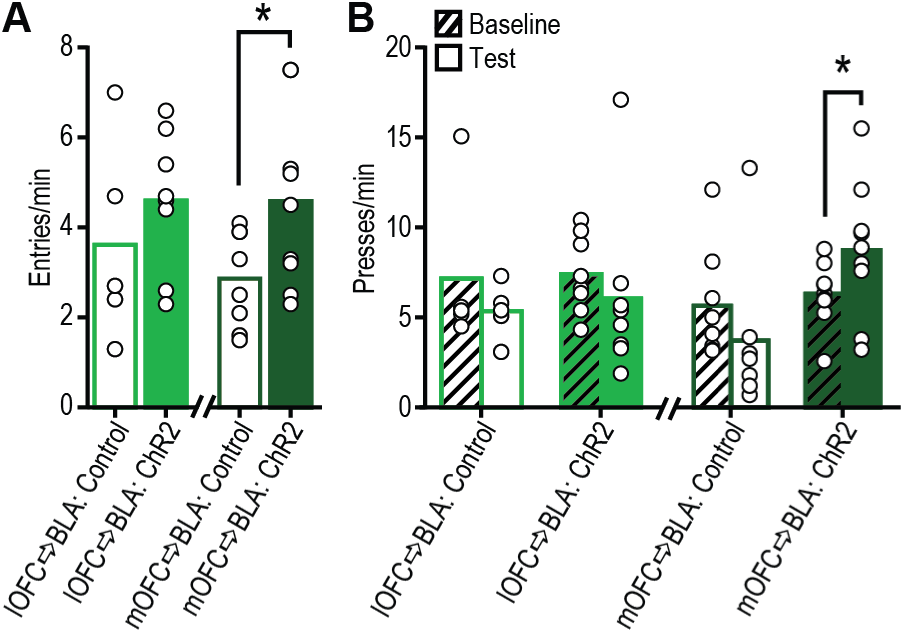
Effect of optical stimulation of lOFC or mOFC terminals in the BLA on reward checking and reward seeking - raw entry/press rates. Following non-contingent sucrose exposure in moderate hunger (8-hr food deprived) state, rats received optical stimulation of lOFC or mOFC terminals in BLA during a non-reinforced, lever-pressing probe test in that moderate hunger state. (**a**) Food-port entry rate (entries/min; lOFC→BLA: *t*_11_=0.94, *P*=0.37; mOFC→BLA: *t*_15_=2.20, *P*=0.04) and (**b**) reward-seeking press rate (seeking presses/min) during baseline and the 8-hr food-deprived non-reinforced lever-pressing probe test with optical stimulation of lOFC (Control, *N*=5; ChR2, *N*=8; Test: *F*_1,11_=1.68, *P*=0.22; Group: *F*_1,11_=0.09, *P*=0.77; Test × Group: *F*_1,11_=0.03, *P*=0.86) or mOFC terminals in BLA (Control, *N*=8; ChR2, *N*=9; Test: *F*_1,15_=0.12, *P*=0.73; Group: *F*_1,15_=3.96, *P*=0.075; Test × Group: *F*_1,15_=9.74, *P*=0.007). \**P*<0.05, between groups (entries) or relative to baseline (presses). Data presented as mean + scatter.

**Supplement Figure 17.**
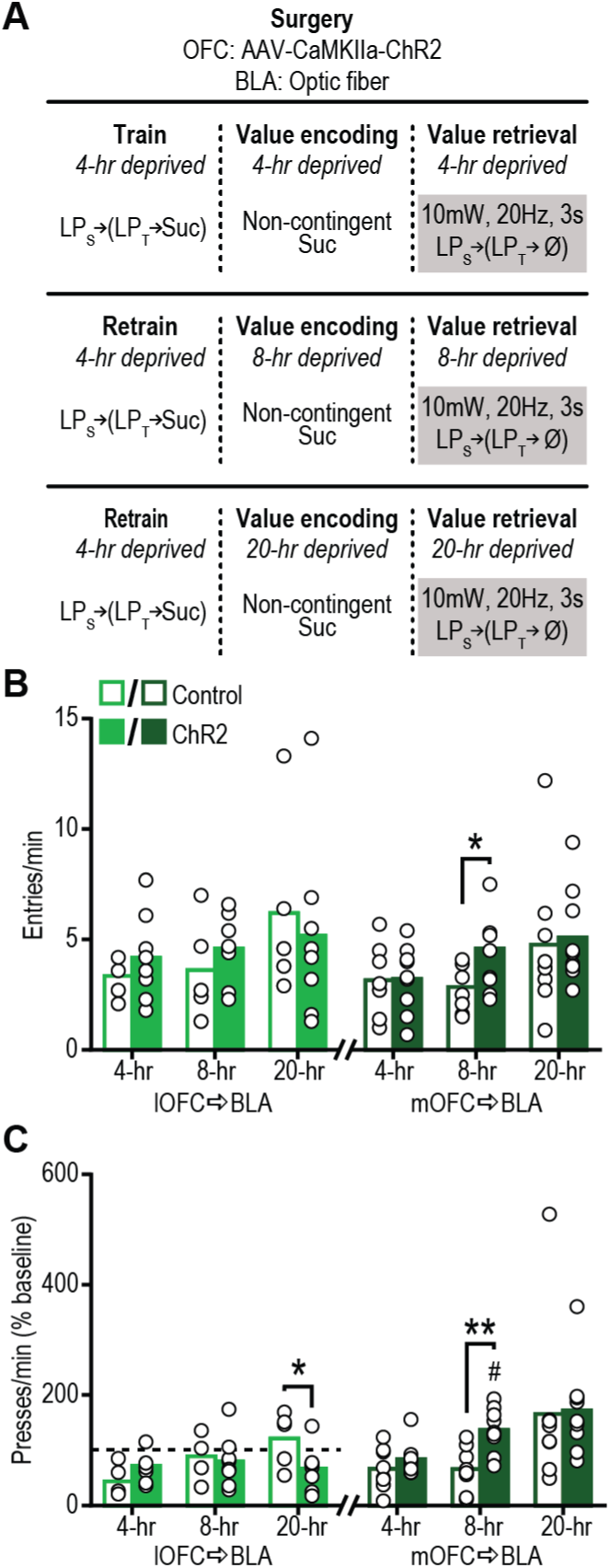
Activation of mOFC, but not lOFC, terminals in the BLA during reward-seeking tests is only sufficient to enhance reward seeking in moderate deprivation state. (**a**) Procedure schematic. Rats received 3 test sets in which they first received non-contingent re-exposure to the sucrose and then, the next day, received a non-reinforced, lever-pressing probe test in the same deprivation state. Light (473 nm, 10mW, 20Hz, 3 s, once/min) was delivered during the lever-pressing test. The control groups consisted of half eYFP + 473 nm and half ChR2 + 589 nm light delivery. Rats were tested at escalating food-deprivation levels (control, familiar 4-hr food deprived state, subthreshold 8-hr food-deprived state, and hungry 20-hr food-deprived state). (**b**) Food-port entry rate (entries/min; lOFC→BLA: Group: *F*_1,11_=0.07, *P*=0.79; Deprivation: *F*_2,22_=1.82, *P*=0.19; Group × Deprivation: *F*_2,22_=0.54, *P*=0.58; mOFC→BLA: Group: *F*_1,15_=0.99, *P*=0.34; Deprivation: *F*_2,30_=4.19, *P*=0.03; Group × Deprivation: *F*_2,30_=1.07, *P*=0.36) during the lever-pressing test. (**c**) Reward-seeking press rate (seeking presses/min), relative to baseline press rate (dashed line), during the probe test with optical activation of lOFC (lOFC→BLA: Group: *F*_1,11_=0.46, *P*=0.51; Deprivation: *F*_2,22_=4.68, *P*=0.02; Group × Deprivation: *F*_2,22_=5.80, *P*=0.009) or mOFC (mOFC→BLA: Group: *F*_1,15_=1.83, *P*=0.20; deprivation: *F*_2,30_=7.81, *P*=0.002; Group × deprivation: *F*_2,30_=0.99, *P*=0.38) terminals in the BLA across escalating food deprivation states. Stimulation of lOFC terminals in BLA was found to disrupt value guided reward seeking following incentive learning. We think this is because stimulating inputs that are not necessary for value retrieval provided an reward value signal unlinked to reward or action, which could have resulted in a contingency degradation-like effect. Planned comparisons: \**P*<0.05, \*\**P*<0.01, between groups. Data presented as mean + scatter. Optical stimulation of mOFC→BLA projections only enhanced reward-seeking activity following subthreshold incentive learning.

**Supplement Figure 18.**
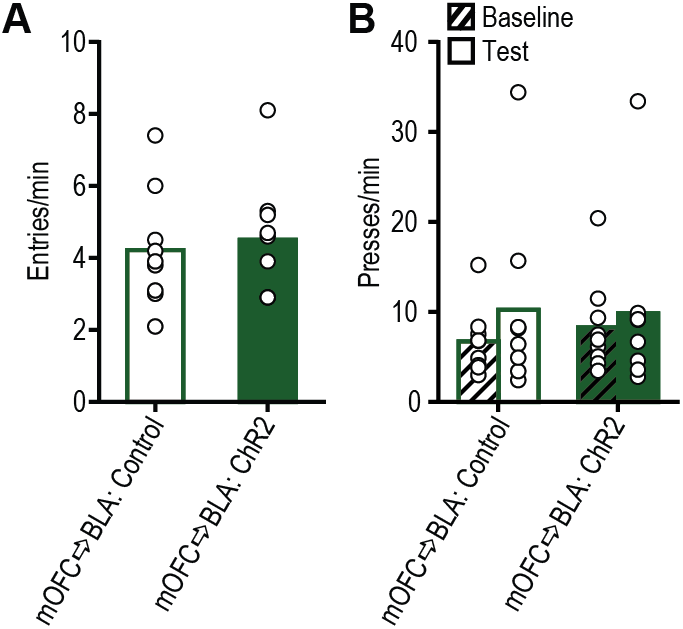
Effect of optical stimulation of mOFC terminals in the BLA on reward checking and reward seeking (no incentive learning) - raw entry/press rates. Following non-contingent sucrose exposure in sated (4-hr food deprived) state, rats received optical stimulation of mOFC terminals in BLA during a non-reinforced, lever-pressing probe test in the moderate (8-hr food deprived) hunger state. (**a**) Food-port entry rate (entries/min) (*t*_8_=0.47, *p*=0.65) and (**b**) reward-seeking press rate (seeking presses/min) during baseline and the 8-hr food-deprived non-reinforced lever-pressing probe test with optical stimulation of mOFC terminals in BLA. Within-subject control consisted of 589 nm light delivery (test order counterbalanced). mOFC→BLA, *N*=9; Test: *F*_1,8_=1.98, *P*=0.20; Group: *F*_1,8_=0.66, *P*=0.44; Test × Group: *F*_1,8_=0.2.29, *P*=0.17. Data presented as mean + scatter.

